# Artificial Tactile Stimulation Provides Haptic Cuing in Force Field Adaptation

**DOI:** 10.1101/2023.07.08.548191

**Authors:** Chen Avraham, Guy Avraham, Ilana Nisky

## Abstract

When interacting with objects with unfamiliar dynamics, the sensorimotor system uses haptic information to develop internal representations of the new dynamics. These representations are subsequently used to manipulate the objects by applying predictive forces that comply with the mechanical properties of the objects. In a recent study (Farajian et al. 2020), we showed that when participants evaluated the stiffness of elastic objects, adding artificial tactile stimulation created an illusion of higher stiffness, increasing the grip force control used to interact with the object. Here, we took a step further in understanding how kinesthetic and tactile information is integrated into the control of objects. Specifically, we examined how added skin stretch influenced the *learning* of novel forces. We found that the extent of force compensation that the participants exhibited depended on the direction of the artificial skin stretch applied simultaneously with the force; learning was enhanced when the skin was stretched in the opposite direction to the external force and diminished when the skin was stretched in the same direction. Strikingly, when the skin stretch stimulation was delivered during probe trials in which the force perturbation was absent, the behavior pattern was flipped, with an increase in force compensation for the same-direction skin stretch stimulation and vice versa. Modeling suggests that these results reflect a unique effect of tactile stimulation during the learning of novel forces; rather than becoming integrated with the dynamic information, it is used by the sensorimotor system as a guidance cue, possibly through explicit mechanisms, providing information on the way to compensate for the forces and optimize movements. We believe that these findings propose a novel instructive role of tactile stimulation during interaction with a dynamic object. This provides a significant potential to leverage these effects in the development of devices aiming to assist and guide users in many human-in-the-loop applications, such as rehabilitation and surgical robotics.

## Introduction

In a competitive rowing tournament, a rower has to have precise control of the oar as she pulls the handle strongly to move the blade against the resisting water. This is one of many examples in which the sensorimotor system gathers multiple types of sensory signals to manipulate objects in a constantly changing environment (J Randall Flanagan et al. 2006; Johansson et al. 2009). Within the haptic modality, this entails the integration of two main sensory signals – kinesthetic and tactile. Kinesthetic information is sensed through mechanoreceptors in the muscles and tendons that capture changes in their respective length and force, while tactile information is sensed by mechanoreceptors in the skin that respond to skin deformations caused by stretch, vibration, pressure, and stroking (Kandel et al. 2000). We recently studied the contribution of the two signals in the perception of stiffness (Farajian et al. 2020; Farajian et al. 2021). Other studies focused on their contribution to the perception of fingertip position (Toma et al. 2019) and object orientation (Frisoli et al. 2011). However, how these signals are integrated during adaptation to new environmental dynamics is largely unknown.

Sensorimotor adaptation allows for maintaining adequate motor performance despite environmental changes. A traditional paradigm for studying sensorimotor adaptation examines how humans adjust their movements and forces in the presence of force perturbations applied by a hand-held robotic device (Conditt et al. 1997; R. Shadmehr et al. 1994; Scheidt et al. 2000; Yousif et al. 2012; Gonzalez Castro et al. 2014; Schween et al. 2020). When exposed to a perturbation, the participants adapt by producing manipulation forces that compensate for the perturbing forces and reduce trajectory errors (G. Avraham et al. 2017; R. Shadmehr et al. 1994; Sing et al. 2009; Thoroughman et al. 2000). In addition to changes in the manipulation forces, adaptation to the perturbation is also associated with changes in the grip forces that are used to prevent slippage of the hand-held device (Hadjiosif et al. 2015; Gordon et al. 1993; Witney et al. 2004; Dafotakis et al. 2008). These changes in the grip force consist of two main components: 1) an overall increase in the baseline (initial) grip force that is non-specific to the perturbing force (Nowak et al. 2002), and 2) the modulation of the grip force throughout the movement with dynamics specific to the perturbing force.

In a recent study, we examined how artificial tactile information changes the manipulation and grip forces during adaptation to force perturbations (C. Avraham et al. 2020). Participants held the handle of a robotic device that applied forces perpendicular to the direction of the movement. To engage the tactile sensors at the gripping fingers, the participants held the device in a precision grip with their fingers placed on rough surfaces and positioned such that the skin of both fingers was stretched due to the applied force (Fig. 1). Additionally, the setup allowed for applying additional (artificial) stretch in the same or opposite direction to that of the perturbing load force. We hypothesized that stretching the skin in the direction of the load force would cause an illusion of a larger force, thus enhancing the adaptive response. Surprisingly, we found an opposite effect: skin stretch in the same direction as the force field attenuated the compensatory response, whereas skin stretch in the opposite direction to the force field increased the adaptive response. When examining the grip forces, we found an increase in both the predictive and reactive components of the grip force for the group that experienced the skin stretch in the same direction of the force field. Conversely, no effect was observed for the group that experienced the skin stretch in the opposite direction.

**Figure 1:**
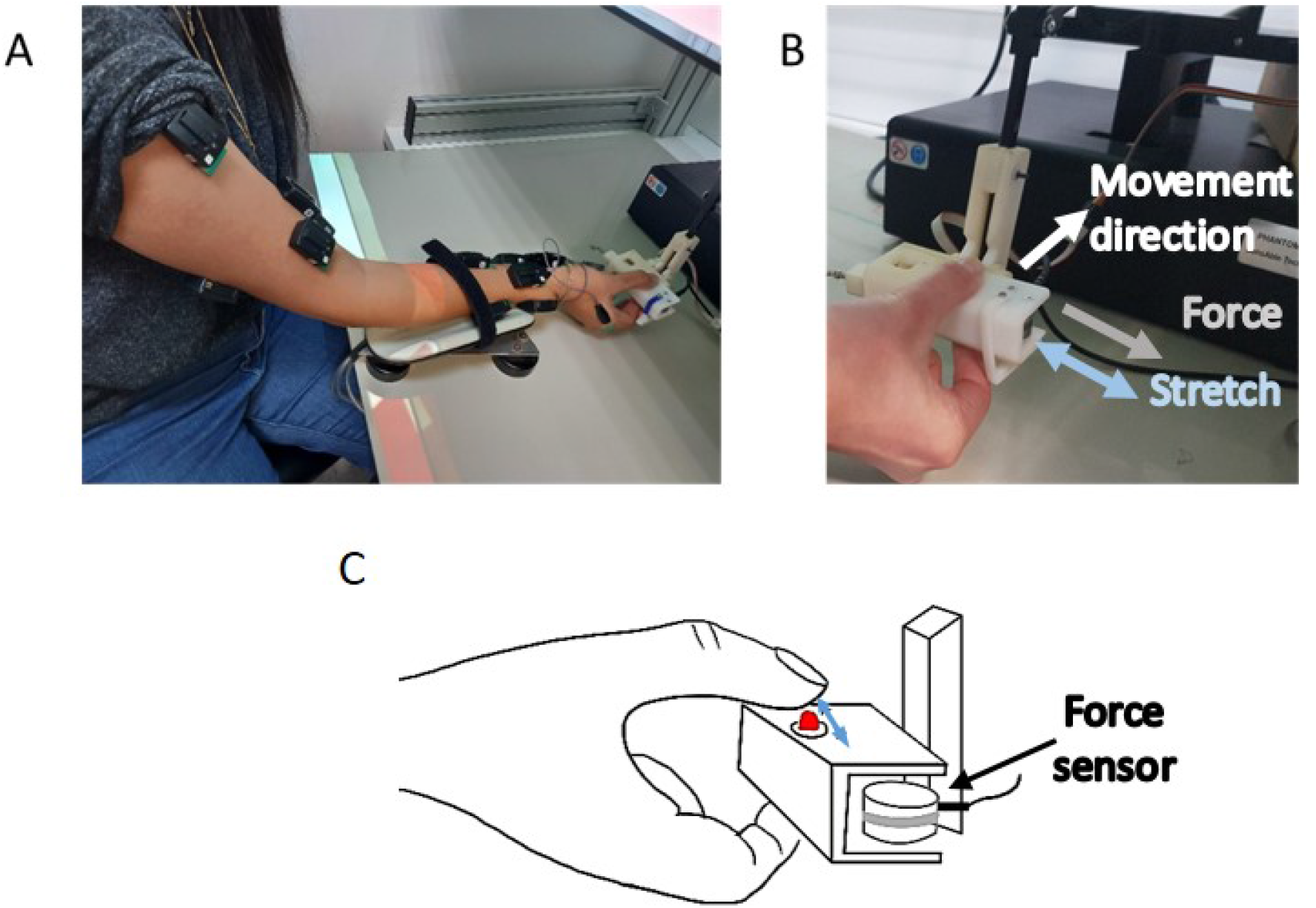
Experimental setup. **(A)** Participants were seated before a screen while holding a skin stretch device attached to a haptic device. Participant’s arm was supported by an air sled to constrain movements to the horizontal plane. Eleven sensors were placed on the right arm to record the EMG signal. **(B)** The skin stretch device. The thumb and index finger were positioned on moving tactors. The grip force was recorded by a force sensor located between the fingers. **(C)** An illustration of the tactor movement direction and location with respect to the participant’s finger. The participant was in contact with the tactor, and the friction between the skin and the coarse surface of the tactor induced stretching of the skin.

Based on the surprising effects of artificial tactile stimulation on the manipulation and grip forces, we propose two hypotheses on how artificial tactile stimulation is integrated with force information during adaptation. The first hypothesis assumes different mechanisms for updating the manipulation and grip forces. For controlling the manipulation forces, the tactile stimulation is interpreted as a guidance cue for the required manipulation forces’ direction, which will compensate for the force field. Accordingly, when the skin stretch is applied in the same direction as the force field, the tactile stimulation will be in the opposite direction to the required manipulation forces. Thus, the stimulation will interfere with adapting to the perturbation. When the skin stretch is applied in the opposite direction to the force field, it will cue the direction and magnitude of the required manipulation forces, assisting the participants in understanding the correct direction for the compensation forces. Independently, the grip force is updated through a slippage sensation caused by tactile stimulation; when the skin stretch is applied in the same direction as the force, it will increase the slippage sensation, thereby increasing the grip force. The second hypothesis stipulates that the manipulation and grip forces are intertwined, such that the additional tactile signal affects both of them through the same mechanism. Specifically, tactile stimulation alters the sensed forces but through a different mechanism than we originally predicted (C. Avraham et al. 2020). Accordingly, the stretch affects the force error felt by the participants during the movement through a slippage sensation that alters the arm co-contraction. Namely, skin stretch in the same direction as the force field causes an increased slippage sensation that increases the arm co-contraction and grip forces. This causes the participant to experience a lower force error than actually produced during the movement. On the other hand, skin stretch in the opposite direction to the force field will decrease the slippage sensation, thereby decreasing the arm co-contraction and increasing the perceived force error.

To differentiate between these two hypotheses, we formulated mathematical equations for the two possible integration models. Thereafter, we designed two experiments. In experiment 1, we aimed to verify the attenuating effect of the artificial tactile information presented concurrently with the force field by exploiting a unique within-participant design. In this paradigm, each participant was exposed to three gains of artificial tactile stimulation – positive (in the force field direction), negative (the opposite direction), and none (no artificial stimulation). In experiment 2, we applied the artificial skin stretch during force channel trials, such that the artificial tactile stimulation was separated from the force field. The results of the experiments showed that tactile stimulation had opposite effects on the manipulation forces when applied together or separately from the force field. In contrast, we observed a similar effect on the grip forces. Therefore, we conclude that tactile stimulation impacts manipulation and grip force control differently. Together with the modeling work, we conclude that in adaptation to perturbing force fields, artificial tactile stimulation does not alter the perceived force information, and, instead, is likely interpreted as a guidance cue directly affecting the motor output.

## Methods

### Computational Models

To understand the effect of artificial tactile information on force field adaptation, we implemented two state-space models. In our previous work, we saw that artificial tactile stimulation had a surprising effect on the control of manipulation and grip forces. Specifically, the skin stretch in the opposite direction to the force field caused participants to apply manipulation forces more similar to the external load forces. In addition, skin stretch in the force’s direction caused participants to use more grip force throughout the adaptation due to the increased predictive and reactive components of the grip force modulation with load force.

To explain the effect on the manipulation forces, we proposed two models for the effect of artificial tactile information on the adaptation process. In both models, we added an internal state which is updated in each trial based on the error sensed by the tactile modality in the previous trial. However, they differ in how this tactile error affects the manipulation forces. The first model assumes that the skin stretch is used as a guidance cue, “instructing” the participants on the proper direction of the manipulation forces they should apply to compensate for the perturbing forces. This is implemented by having the tactile stimulation affect only the final motor movement rather than the trial-to-trial learning process. The second model assumes that the skin stretch error is integrated with the force error to update the trial-to-trial learning process. This hypothesis assumes that tactile and kinesthetic cues are integrated to estimate the applied force, as was previously proposed for the effect of skin stretch on the perception of stiffness (Zhan Fan Quek et al. 2014). For each model, we estimated the parameters (retention factor A, learning rate B, and the weighting parameter *α* that are marked in bold in equations 1, 2, 4, 10, and 12 below) that yield the closest results to the data from our previous study (C. Avraham et al. 2020), using the function `lsqcurvefit` in Matlab. Then, we examined the validity of our models in the results of the two experiments in the current study.

In all the equations above, *A* is the retention factor, *B* is the learning rate (or error sensitivity). In both models, the internal state in each trial (*x*^(*n*)) is comprised of fast and slow processes acting in parallel (Smith et al. 2006), such that each internal state depends on the output and the error at the previous trial:

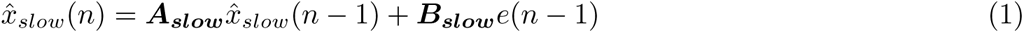

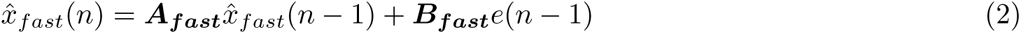

The internal state is the sum of the two processes:

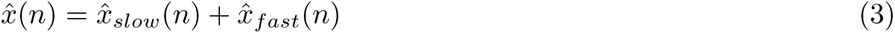

Following exposure to novel dynamics, an error is sensed between the force field and the internal state:

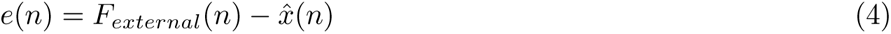

*F_external_*(*n*) is the external force, such that:

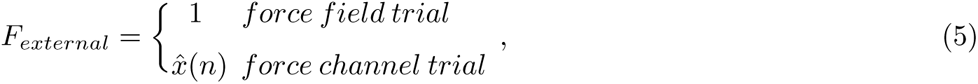

as defined in (Smith et al. 2006).

To assess the contribution of the artificial tactile information, we added a tactile state and a tactile error:

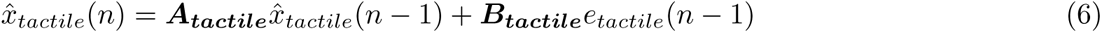

The tactile modality is updated according to the error between the estimated tactile sensation *x*^*_tactile_* and the actual tactile stimulation *F_tactile_*(we note that, in this case, the participant is unable to cancel the applied stretch):

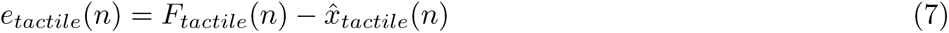

Similarly to the external force, *F_tactile_*(*n*) is the artificial tactile stimulation defined as

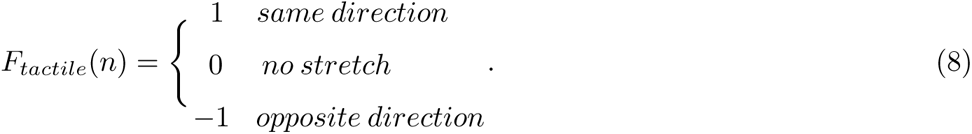

We note that in all conditions (including no stretch), the skin experiences a natural stretch due to the force field, which is not addressed here for simplicity (see discussion for more details). In addition, we assumed a single tactile state, but the model could be easily extended to include slow and fast tactile states.

Essentially, the models differ in the way the motor output is calculated and the error in each trial. In both models, the final motor output depends on the internal state (*x*^(*n*)) and internal noise (*η* ∼ *N* (0, 1)) that represents the variability in the motor system (Reza Shadmehr et al. 2008):

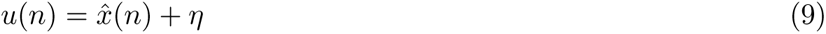

#### Model 1-Haptic cuing

This model suggests that tactile stimulation is interpreted as a signal that guides the participant on the desired motor output. Therefore, the tactile state directly affects the final motor output in each trial:

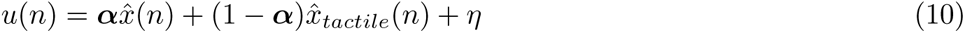

where *α* is a weighting parameter between the internal and tactile states.

#### Model 2-Error integration

This model suggests that tactile stimulation affects the perceived forces, causing the participants’ sensorimotor system to misinterpret the magnitude of external force applied to them. We implemented this idea by considering the experienced error as a combination of the force and tactile errors. The error in each trial is calculated similarly to the previous model, except being signified as *e_force_*(*n*):

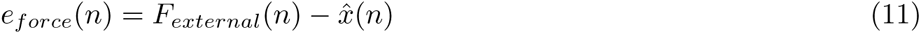

The overall error perceived by the participants and used to calculate the internal states at the next trial is then affected by the tactile and force errors:

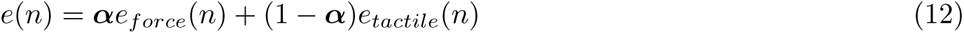

Similarly to the previous model, *α* is a weighting parameter between the force and tactile errors.

**Table 1:** Models summary

|  | Model 1- Haptic cuing | Model 2- Error integration |
| --- | --- | --- |
| **Motor output** | $u(n) = \alpha \hat{x}(n) + (1 - \alpha) \hat{x}_{tactile}(n) + \eta$ | $u(n) = \hat{x}(n) + \eta$ |
| **Force error** | $e(n) = F_{external}(n) - \hat{x}(n)$ | $e(n) = \alpha [F_{external}(n) - \hat{x}(n)] + (1 - \alpha) [F_{tactile}(n) - \hat{x}_{tactile}(n)]$ |

### Participants and setup

Seventy-one right-handed participants (33 females and 38 males, aged 22-29) participated in the experiments. The participants were naive to the purpose of the investigation and were paid to participate. The experiment was approved by the Human Participants Research Committee of Ben-Gurion University of the Negev, Beer-Sheva, Israel.

The experiment was performed with a PHANTOM^®^ Premium 1.5 haptic device, which recorded kinematic (position and velocity) and dynamic (manipulation forces) data at a 200Hz sampling rate, applying forces on the participants’ arm. The device was located beneath a white horizontal surface with a projector displaying the visual stimuli. On top of the participants’ right arm, we attached EMG sensors (Delsys Trigno™ Wireless System) that recorded muscle activity: five on the upper arm, three on the forearm, and three on the palm. We used an air-sled wrist supporter, which allowed planar movements with minimal friction. The participants wore noise-canceling headphones to mask the motor and wrist supporter noises. A black fabric sheet covered their upper body to prevent the visual image of the hand (Fig. 1A).

### Skin stretch device

We used a custom-designed 3D-printed device (a modified version of the one used in (Farajian et al. 2020)) attached to the PHANTOM haptic device with a single rotational degree of freedom. The skin stretch module consisted of a rail that enabled one degree of freedom of movement for the tactors. The tactors were located in the center of an aperture (one on each side of the device), such that the upper surface of the tactor was slightly above the height of the device’s flat surface. The aperture size restricted the movement of the tactors to a maximum translation of 2.5 mm to each side, left and right. Participants held the device using their thumb and index fingers, where the fingers were in contact with the device’s surface around the aperture (static, natural tactile sensation) and the upper surface of the tactors (dynamic, artificial tactile sensation) (Fig. 1C). With this configuration, the stretch was applied in the same plane (horizontal) as the movement direction and the applied force field and perpendicular to the movement axis (Fig. 1B). For more details regarding the device’s configuration, see (Farajian et al. 2020).

The actuation was obtained via a DC motor, a spur gearhead, and an encoder (discussed elaborately in (C. Avraham et al. 2020)). Grip forces were recorded through a force sensor attached to the right end of the device. The grip force between the thumb and index fingers was transmitted to the sensor by a “door” that pressed the force sensor with an amount relative to the grip force applied on the other side, as in (C. Avraham et al. 2020; Farajian et al. 2020).

### Protocol

During both experiments, the participants performed reaching movements in a sagittal plane from a start position (white circle, 1 mm radius) toward a target (Black circle, 1 mm radius) located 10cm away (Fig. 2). The movement of the participants controlled a cursor (yellow circle, 0.5 mm radius) displayed on top of a screen. A trial was initiated when the cursor was at the start position for 1.5 sec. Subsequently, an imperative cue was initiated when the start position turned green, signaling the participant to perform a reaching movement toward the target. The trial ended when the movement velocity was less than 0.05 cm/sec. Next, we presented feedback regarding the speed of the movement: “Move slower” if the movement duration was lower than 0.4 sec, “Move faster” if the movement duration was higher than 0.6 sec, or “Exact” otherwise.

**Figure 2:**
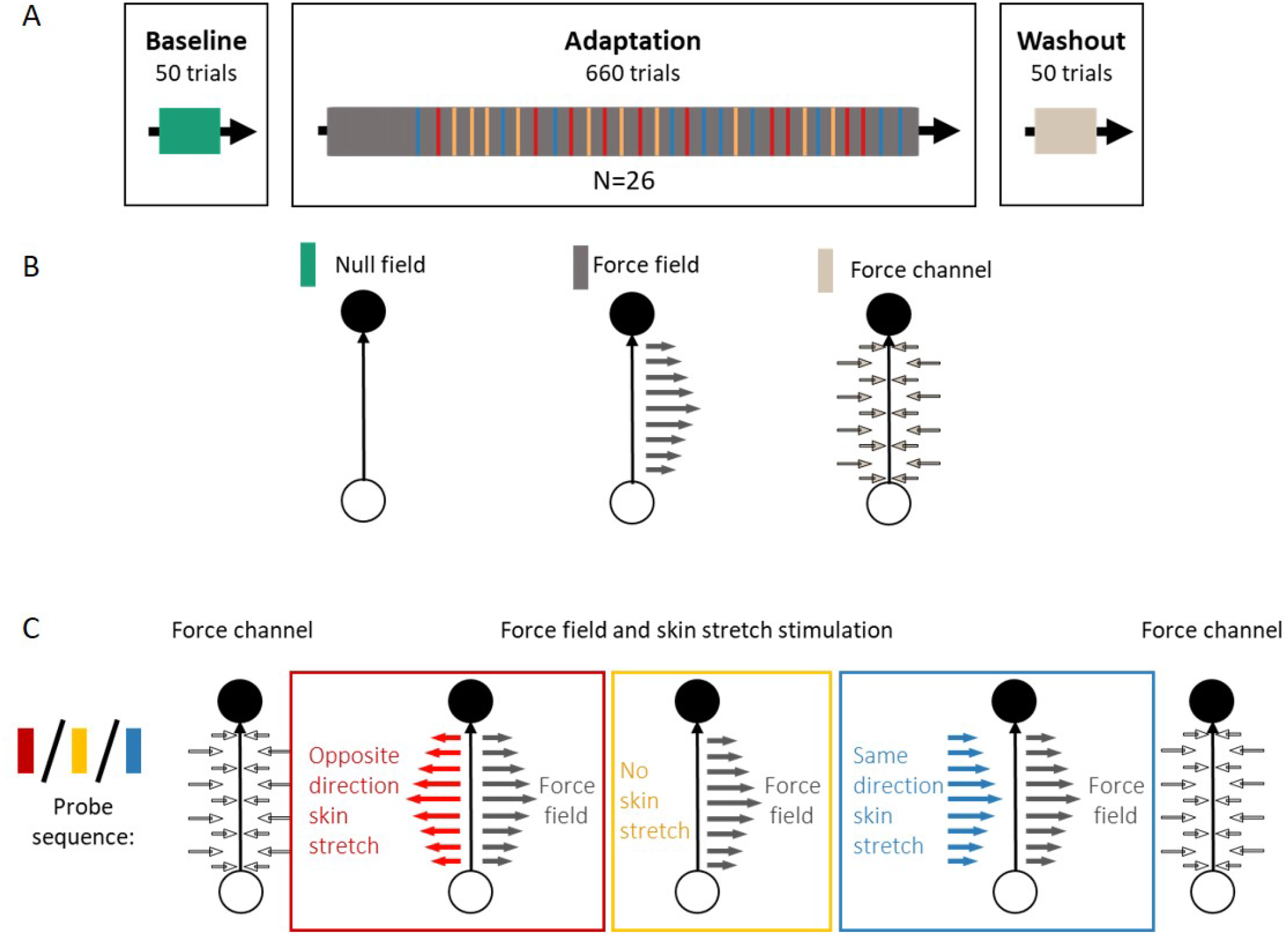
Experiment 1-protocol. **(A)** The experiment consisted of three blocks: Baseline, Adaptation, and Washout. In the Baseline block, we presented null-field trials (green bar). In the Adaptation block, we presented force field trials (gray bar) and probe sequences (red/yellow/blue). The Washout block included only force channel trials (beige). **(B)** In null field trials (green bar), the participants performed reaching movements from a start position (white circle) toward a target (black circle) without experiencing any perturbation. In force field trials (gray bar), participants were exposed to a velocity-dependent force field. In force channel trials (beige), we applied virtual walls that constrained movements to a straight path, allowing us to measure compensatory manipulation forces. **(C)** During Adaptation, we presented probe sequences consisting of trial triplets: force channel – force field with skin stretch (red/blue) or without skin stretch (yellow) – force channel. The skin stretch was presented in the force field direction (blue) or the opposite direction (red).

Each experiment consisted of three blocks: Baseline, Adaptation, and Washout. Throughout the experiment, the participants experienced three types of trials: null field, force field, and force channel. During force field trials, we applied velocity-dependent perturbing forces by the haptic device:

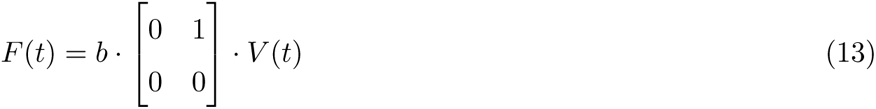

where 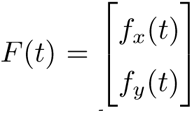 is the force field, *b* = 10[*N* ・ *s*/*m*] is the velocity gain, and 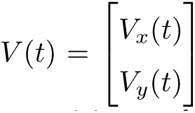 is the velocity.

Accordingly, the force was perpendicular to the straight line connecting the start position and target. In null and force field trials, the participants were able to move in the horizontal plane. During force channel trials, we presented virtual walls with a viscoelastic force field, inhibiting any lateral forces and thereby imposing a straight trajectory:

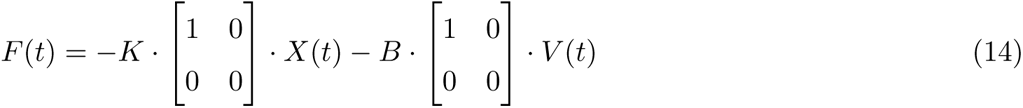

where *K* = 500[*N/m*] is the walls’ stiffness, *B* = 2[*N · s/m*] is the viscosity, *X*(*t*) is the position, and *V* (*t*) is the velocity. The forces applied by the participants towards the virtual walls were considered as proxies to the manipulation forces they applied to resist the forces in force field trials (Scheidt et al. 2000; Joiner et al. 2008). To augment the skin’s natural stretch, we applied a velocity-dependent skin stretch stimulation by moving tactors attached to the skin of the thumb and index finger:

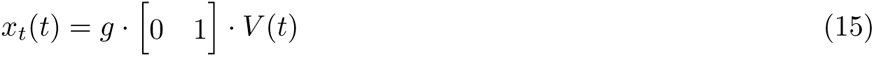

where *g* = {*−*100, 0, 100} [*mm · s/cm*] is the tactors’ displacement gain, and *V* (*t*) is the velocity.

#### Experiment 1

26 participants participated in this experiment (12 females and 14 males). The Baseline block consisted of 50 null-field trials. The purpose of this block was to acquaint participants with the reaching task and establish baseline performance. The adaptation block consisted of force field trials (600 trials). In addition, 10% of the trials during adaptation (60 trials) were force channel trials. The trials were placed within the block, forming *probe sequences* of trial triplets consisting of force channel – force field – force channel (Fig. 2). The force field trials Within each probe sequence were accompanied by either negative, null, or positive tactile stimulation (*g* = *−*100, *g* = 0, or *g* = 100, respectively). All participants were presented with all three levels of skin stretch to allow for a within-participant comparison. The probe sequences were presented in random and predetermined locations within the adaptation block. However, the following constraints were included: 1) The first probe sequence was presented following at least 100 force field trials. This constraint ensured the tactile stimulation effect was assessed only after participants formed a reliable representation of the force field. Our previous study showed that with the current setup, 100 trials were sufficient for obtaining full-force compensation (C. Avraham et al. 2020). 2) participants experienced at least 12 force field trials between probe sequences to minimize the potential effects of forgetting during the force channel trials (Scheidt et al. 2000). The washout block included 50 force channel trials. It was added to maintain a similar experimental design to our previous experiments (C. Avraham et al. 2020)

#### Experiment 2

In Experiment 2, we aimed to investigate whether the artificial tactile information would affect the adaptation when presented in trials with no kinematics errors, i.e., in the absence of both the kinesthetic and natural tactile information obtained during force field exposure. Therefore, we decoupled the artificial stretch from the force field and presented it only during force channel trials (Fig. 3A-C). 45 participants (21 females and 24 males) took part in this experiment designed for a between-participants comparison. The Baseline block included 50 trials of null field reaching movements. The Adaptation block included 300 trials, divided into 260 force field trials and 40 force channel trials. In half of the force channel trials, artificial skin stretch was applied, for which we considered three levels of stretch (presented for three different groups of participants): (1) skin stretch in the opposite direction to the force field (*g* = *−*100*, N* = 15), (2) skin stretch in the same direction as the force field (*g* = 100*, N* = 15), and (3) control without skin stretch (*g* = 0*, N* = 15) (Fig. 3C). Participants were assigned randomly to the different stretch-level groups. In the washout block, we conducted 100 trials of null field and force channel without skin stretch.

**Figure 3:**
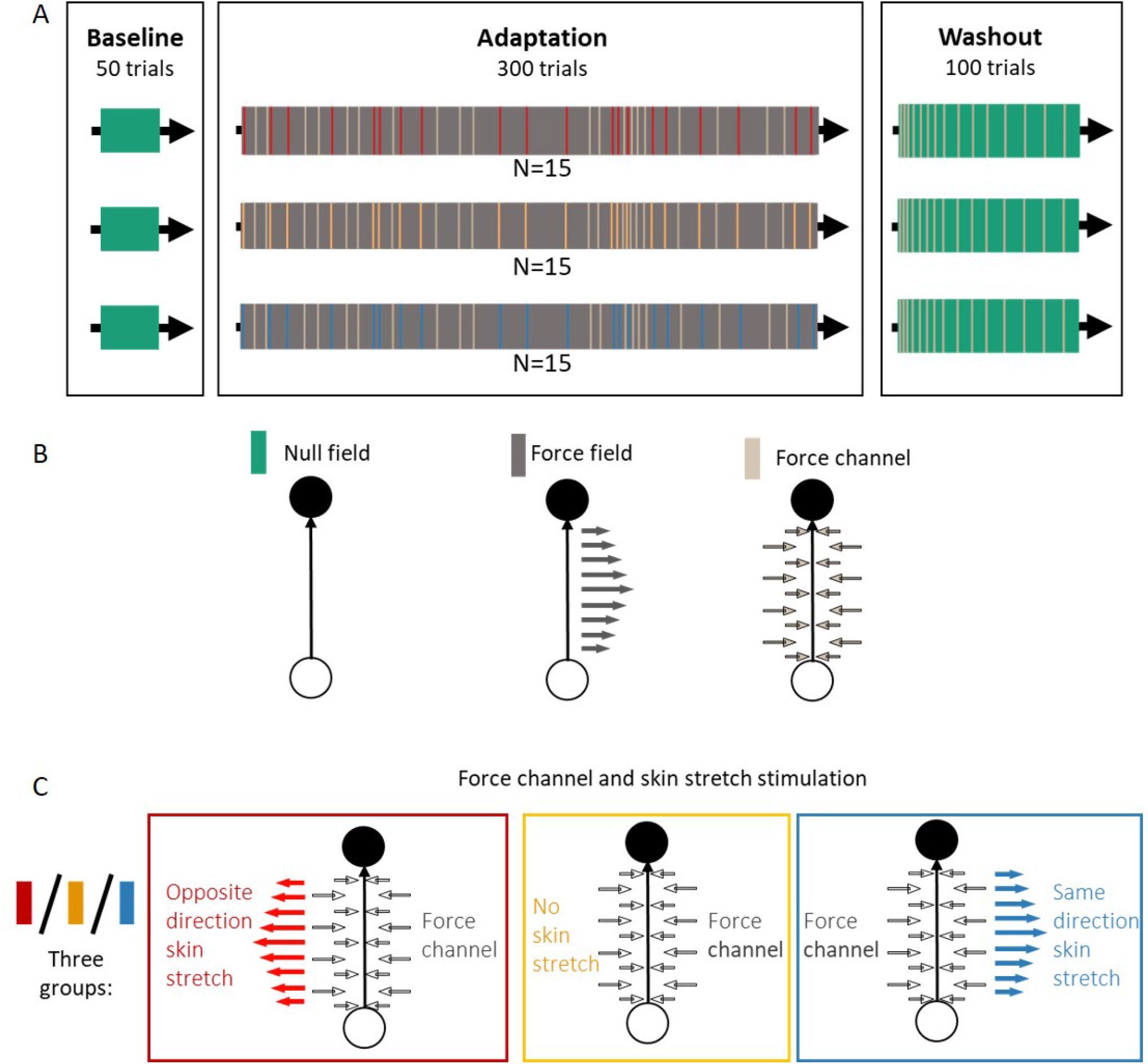
Experiment 2-protocol. Three groups of participants took part in this experiment. **(A)** Three blocks were provided for all groups: Baseline, Adaptation, and Washout. In the Baseline block, we presented null-field trials (green bar). The Adaptation block included force field trials (gray bar) or force channel trials with and without skin stretch (beige/red/yellow/blue). Essentially, there is no difference between beige and yellow bars, as they both represent force channel trials without skin stretch. However, yellow bars represent force channel trials in the control group that were accompanied by skin stretch stimulation for the other two groups, while beige bars represent force channel trials without stimulation for all groups. The Washout block comprised both null field trials and force channel trials without skin stretch. **(B)** In null field trials (green bar), the participants performed a reaching movement from a start position (white circle) toward a target (black circle) without experiencing any perturbation. In force field trials (gray bar), participants were exposed to a velocity-dependent force field. In force channel trials (beige), we applied virtual walls that constrained movements to a straight path. **(C)** During Adaptation, in half of the force channel trials, we presented a velocity-dependent skin stretch that was either in the direction of the force field (blue) or the opposite (red). Each group experienced only one skin stretch stimulation type. The third, control group, did not experience any skin stretch (yellow).

## Data analysis

Data were analyzed offline using a custom-written MATLAB code (The MathWorks, Inc., Natick, MA, USA). We used two main measures for assessing the effect of tactile stimulation on adaptation: directional error and force compensation. In addition, we examined the grip forces and muscle activity to understand further the effect of tactile stimulation on internal representation. The directional error was defined as the maximum perpendicular deviation from the straight line connecting the start position and the target. This measure was obtained from all the trials that did not include force channels.

The adaptation coefficient was calculated as the regression slope between the forces participants applied during force channel trial and the force field applied in the previous trial (with intercept). We interpolated each signal to have a fixed number of samples using the Matlab function `interp1()`, to match the temporal alignment between the two signals. In experiment 1, we were interested in the skin stretch effect on the manipulation forces applied in the force channel trial after exposure to the force field (last trial in the probe sequence), compared to the manipulation forces that were applied in the force channel trial before the exposure (first trial in the probe sequence). Therefore, we calculated the *adaptation difference* for each value of skin stretch gain by subtracting the adaptation coefficient in the trial before the exposure from the coefficient after the exposure.

To assess the ability of the participants to adjust the grip force according to the applied force field and tactile stimulation, we calculated the modulation between the grip force and the load force:

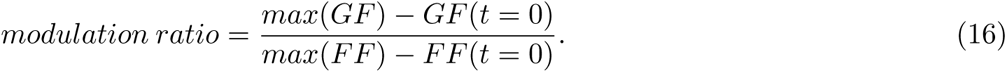

The force-specific modulations typically include a predictive component based on the formed representation of the external forces throughout the experience and a reactive component activated online as soon as the system detects an unexpected change during movement. For the predictive grip force analysis, we were interested in the effect of the single exposure with the force field and artificial skin stretch on the internal representation formed during adaptation. Therefore, the effect on the grip force was assessed by computing the *modulation ratio difference*, as the difference between the measure calculated in the trial wherein the tactile stimulation was abruptly removed (force channel after the exposure) and the previous trial without tactile stimulation (force channel before the exposure). To extract the reactive grip force, we calculated the same measure of *modulation ratio difference*; however, here, we compared a trial where the stretch was surprisingly introduced (force field trial within the probe sequence) to a trial with no skin stretch (force field trial before the probe sequence). This analysis was performed separately for each gain of skin stretch. In experiment 2, we calculated the modulation ratio in force field trials and force channel trials with and without the skin stretch. The predictive and reactive components of the modulation grip force were assessed by comparing the modulation ratio between the two types of force channel trials.

In addition, we calculated the baseline grip force (*GF* (*t* = 0)) to establish how certain the participants were about the external dynamics. In experiment 1, we calculated the *baseline GF difference* similar to the *modulation ratio difference*. In experiment 2, we examined the baseline grip force applied in force field trials and force channel trials with and without the skin stretch.

To examine participants’ muscle activity during the experiment, we recorded the EMG signal from eight muscles in the right arm: the Deltoid Lateral, Biceps Short, Biceps Long, Triceps Long, Triceps Lateral, Flexor Carpi Radialis, Flexor Carpi Ulnaris, and Extensor Carpi Ulnaris. In addition, we recorded the muscle activity of three muscles in the right palm to verify the results of the grip forces: Adductor Pollicis, Adductor Pollicis Brevis, and First Dorsal Interosseus. Data collection was obtained using the Delsys Trigno™ Wireless EMG System. Data preprocessing was performed offline using MATLAB and included band pass filtering (15-400Hz) with a second-order Butterworth filter using the Matlab `filtfilt()` function, sampling the data at 2kHz, amplifying, and full-wave rectifying. To adjust the scale of the signals from the different muscles, we calculated the mean EMG signal for each muscle in the last five trials of the pre-exposure phase and normalized each signal by the corresponding mean. Integrated EMG (iEMG) was calculated over the interval of [100*ms* – 600*ms*], which includes online corrections of muscle activity and reflexes (Milner et al. 2005). Accordingly, an increase in the iEMG measure would indicate an increase in muscle activity and, therefore, in muscle co-contraction (Heald et al. 2018).

### Statistical analysis

For the analysis of experiment 1, we were interested in assessing the effect of the skin stretch gain on the averaged adaptation difference, baseline grip force difference, and modulation ratio difference. Hence, we fitted a one-way repeated measures ANOVA with a within-participant factor of stimulation gain (*g* = *−*100, 0, 100). Significant effects were defined at the *p <* 0.05 level. A post-hoc t-test was performed with a Bonferroni correction when found to have a significant main effect. To analyze the adaptation coefficient at the end of adaptation (5 last probe trials) and the integrated EMG (see Supplementary Information), we examined whether the effect was linearly dependent on the amount of stretch. Therefore, we fitted a linear regression model to the mean values of the adaptation coefficient and integrated EMG as a function of the three gains of skin stretch.

In experiment 2, we considered three groups of participants, so each group was exposed to a different stretch gain. Unlike in experiment 1, here we aimed to assess the stretch effect in the different stages of the adaptation process. To do so, we implemented two statistical tests. The first was a two-way mixed-design ANOVA with a between-participants factor (opposite direction/control/same direction) and a within-participants factor of the stage along the experiment. For the directional error, we examined the difference between four stages (five trials from each stage): the end of the Baseline block (Late Baseline), the beginning and end of the Adaptation block (Early and Late Adaptation, respectively), and the beginning of the Washout block (Early Washout). When analyzing the manipulation and grip force control, we compared the performances at the beginning and end of the Adaptation block (Early Adaptation and Late Adaptation). In this experiment, we added a second statistical analysis, a nonparametric permutation test, which allows the examination of the differences between the groups by considering all adaptation trials without the need to define separate stages arbitrarily (Maris et al. 2007). This analysis detects clusters of interest in the data using an ANOVA test assigning a “score” for each cluster. Namely, for each trial, we assigned an F-value by performing a one-way ANOVA test with one between factor of group (opposite direction/control/same direction). Then, we clustered together consecutive trials with a p-value lower than a threshold (0.05) by summing their F-values. We assessed the significance of these clusters against clusters obtained in 10,000 permuted data sets. In each permutation, participants were randomly assigned to each of the three groups; we performed the same cluster detection as in the original data and saved the highest F-values, generating a distribution of F-values for clusters obtained by chance. Finally, we evaluated how often the “score” of our original cluster of interest was higher than the scores detected in the permutation test and divided it by the number of permutations, providing us with the Monte Carlo p-value. When significant main effects were found (*p <* 0.025), the same analysis was performed within every two groups using a two-sample t-test and compared to an adjusted p-value (*p <* 0.025).

## Results

To better understand the effect of artificial tactile stimulation on force field adaptation, we aimed to computationally model this effect using state space equations. We first constructed the equations along with an additional tactile state, to help us distinguish between the different conditions of tactile stimulation during the experiment. Then, we proposed two possible hypotheses for the effect of artificial tactile stimulation on force field adaptation. To verify our model and distinguish between the possible hypotheses, we performed two experiments. Experiment 1 aimed to replicate our previous findings that showed interesting adaptation effects; for tactile stimulation in the opposite direction to the force field, we observed increased adaptation, whereas decreased adaptation was found for tactile stimulation in the force field direction. To increase sensitivity, we chose a unique within-participant design, where each participant was exposed to all skin stretch gains. In experiment 2, we separated the force field and artificial tactile stimulation to enable us to refute one of the possible hypotheses.

## Modeling

We used state space equations with two implementations that could explain the effect of tactile stimulation on force field adaptation. Exploiting the state space framework, we could examine potential mechanisms for the effect of artificial tactile information on the adaptation process. Mainly, we focused on two mechanisms of haptic cuing and error integration.

In both models, we included seven parameters: the retention factors of the three internal states of slow, fast, and tactile (*A_slow_*, *A_fast_*, and *A_tactile_*, respectively), learning rates (*B_slow_*, *B_fast_*, and *B_tactile_*), and the weighting parameter *α*. For each group with a different skin stretch gain (*g* = *−*100, *g* = 0, and *g* = *−*100), the parameters were estimated according to the adaptation curve received in our previous study (C. Avraham et al. 2020). Accordingly, as in this study, the force field and tactile stimulation were presented together, and the external force *F_external_*(*n*) and artificial tactile stimulation *F_tactile_*(*n*) were correlated (*F_external_*(*n*) = 1 *⇒ F_tactile_*(*n*) = 1*/ −* 1). Subsequently, we used the parameters that yielded the best fit to compute the motor output *u*(*n*) when the tactile stimulation was presented during force channel trials (*F_external_*(*n*) = *x*^(*n*) *⇒ F_tactile_*(*n*) = 1*/ −* 1). For the control group, there was no tactile error, and consequently, the tactile state *x*^*_tactile_* was zero in both conditions.

In the first model, the computation yielded an opposite effect on the manipulation forces between the two conditions in which the skin stretch was presented in force field or force channel trials (Fig. 4, left panel). In contrast, the second model yielded a similar effect in the two conditions, with a higher effect when the stretch was applied in force channel trials (Fig. 4, right panel).

**Figure 4:**
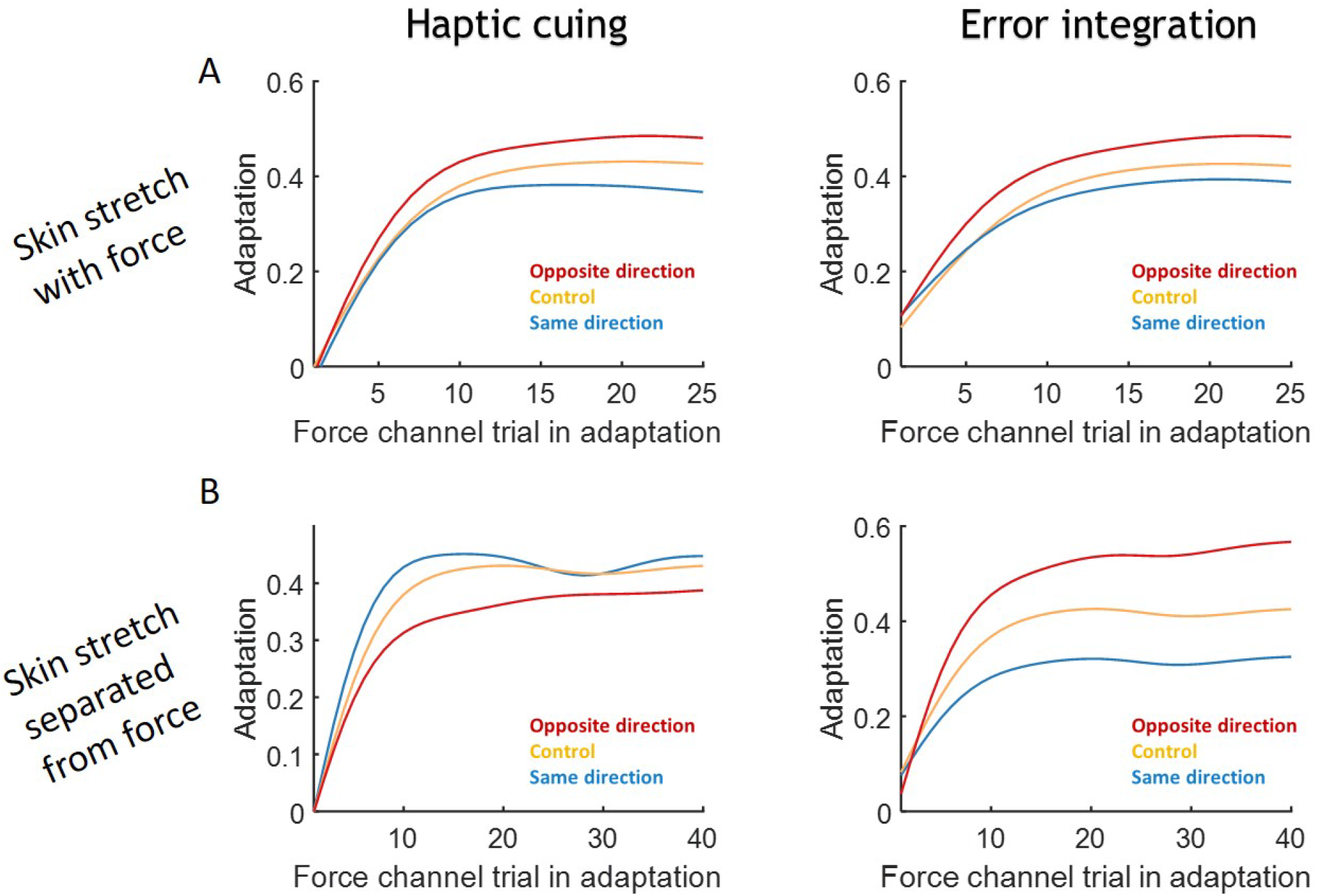
Modeling results. **(A)** Simulation results for the two hypotheses regarding the effect of tactile stimulation on adaptation: a direct effect on the motor output (Haptic cuing, right) or an effect on the perceived force error (Error integration, left). The parameters were fitted according to the results of our previous study (C. Avraham et al. 2020). Results are presented for the three groups of skin stretch in the opposite direction to the force field (red), control (yellow), and the same direction as the force (blue). **(B)** Simulation results for experiment 2 in which the skin stretch was separated from the force field. Colors are marked as in (A).

### Augmented tactile stimulation in the same direction of a perturbing force reduces force compensation

In experiment 1, we considered a within-participant design, wherein each participant was exposed to three skin stretch gains in a sequence of three trials: force channel – force field with or without skin stretch – force channel. This allowed a sensitive assessment of the skin stretch effect on the adaptation process, which would appear when comparing the force channel trials before and after the stimulation and the force field trials with and without the stimulus.

#### Directional error

We first examined the directional error curve to verify that all probe sequences were presented in the phase where participants adapted to the perturbation. When experiencing the force on the first trial, participants underwent a large directional error (Fig. 5). This error decreased with continuous exposure, such that the learning curve reached a value equal to *e^−^*^1^ of the initial error after six reaching attempts. Moreover, after 100 trials in adaptation, the directional error was reduced to an approximately steady-state value. Therefore, we confirmed that we could average all probe sequences from this period.

**Figure 5:**
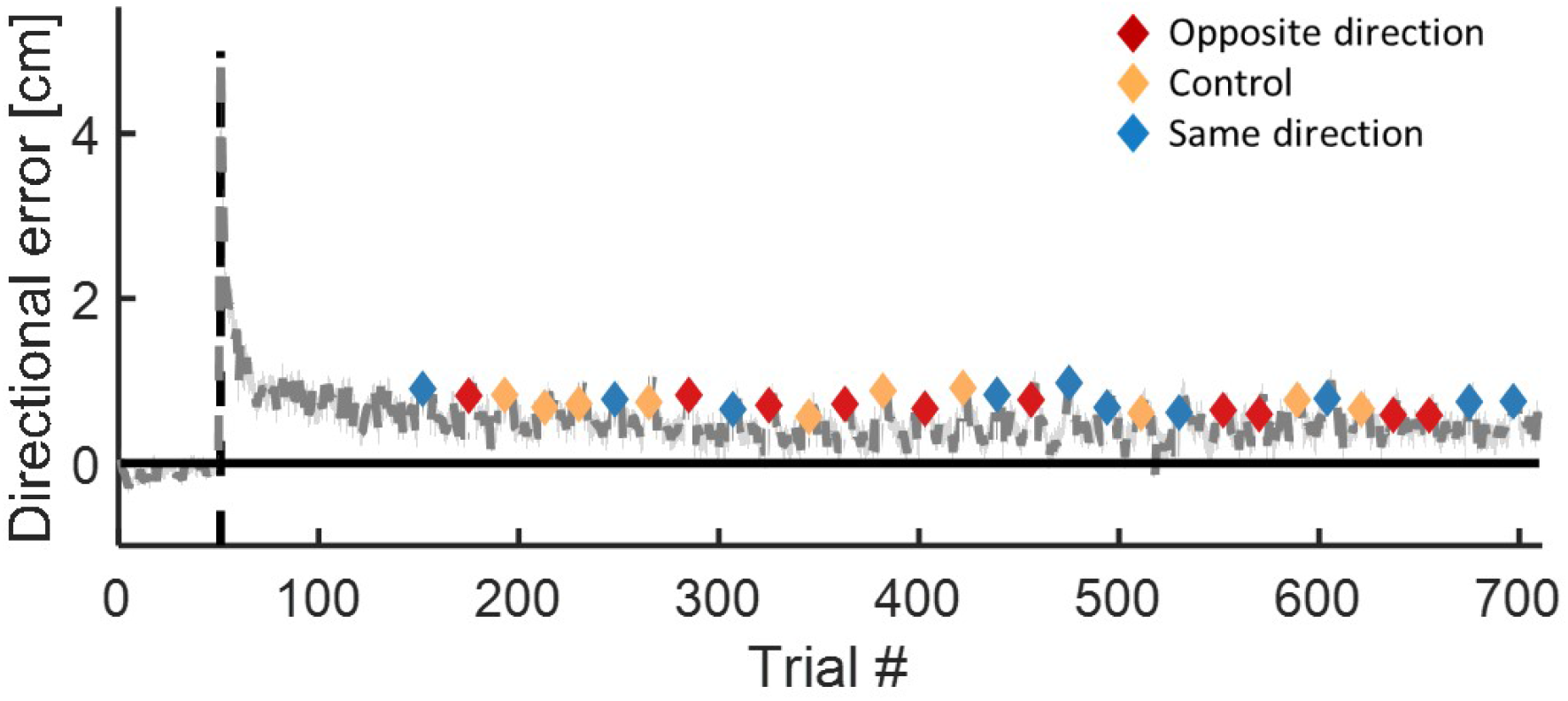
Experiment 1-Directional error. Averaged maximum deviation from a straight line for all participants in the baseline and adaptation blocks. Markers represent the directional error in force field trials within a probe sequence. The dashed black line signifies the beginning of the Adaptation block, and the shaded region is ±SE.

#### Force compensation

To examine the effect of tactile stimulation on adaptation, we calculated the adaptation coefficient in all force channel trials during adaptation (Fig. 6A). Then, we compared the performances in the two force channel trials within a probe sequence. We did so by subtracting the adaptation coefficient at the trial before the tactile stimulation from the adaptation coefficient after the tactile stimulation in all probe sequences during adaptation. Since we verified that all probe sequences were in the steady state phase of adaptation using the directional error, we averaged all coefficients (*mean adaptation difference*) for each stimulation type (Fig. 6B). We found that the skin stretch in the same direction as the force field decreased the adaptation coefficient in the proceeding trial after the exposure. However, this effect was not significant (ANOVA model: *F*_2,50_ = 1.40, *p* = 0.255, regression model: *p* = 0.12). In addition, no effect was observed for the opposite skin stretch direction.

**Figure 6:**
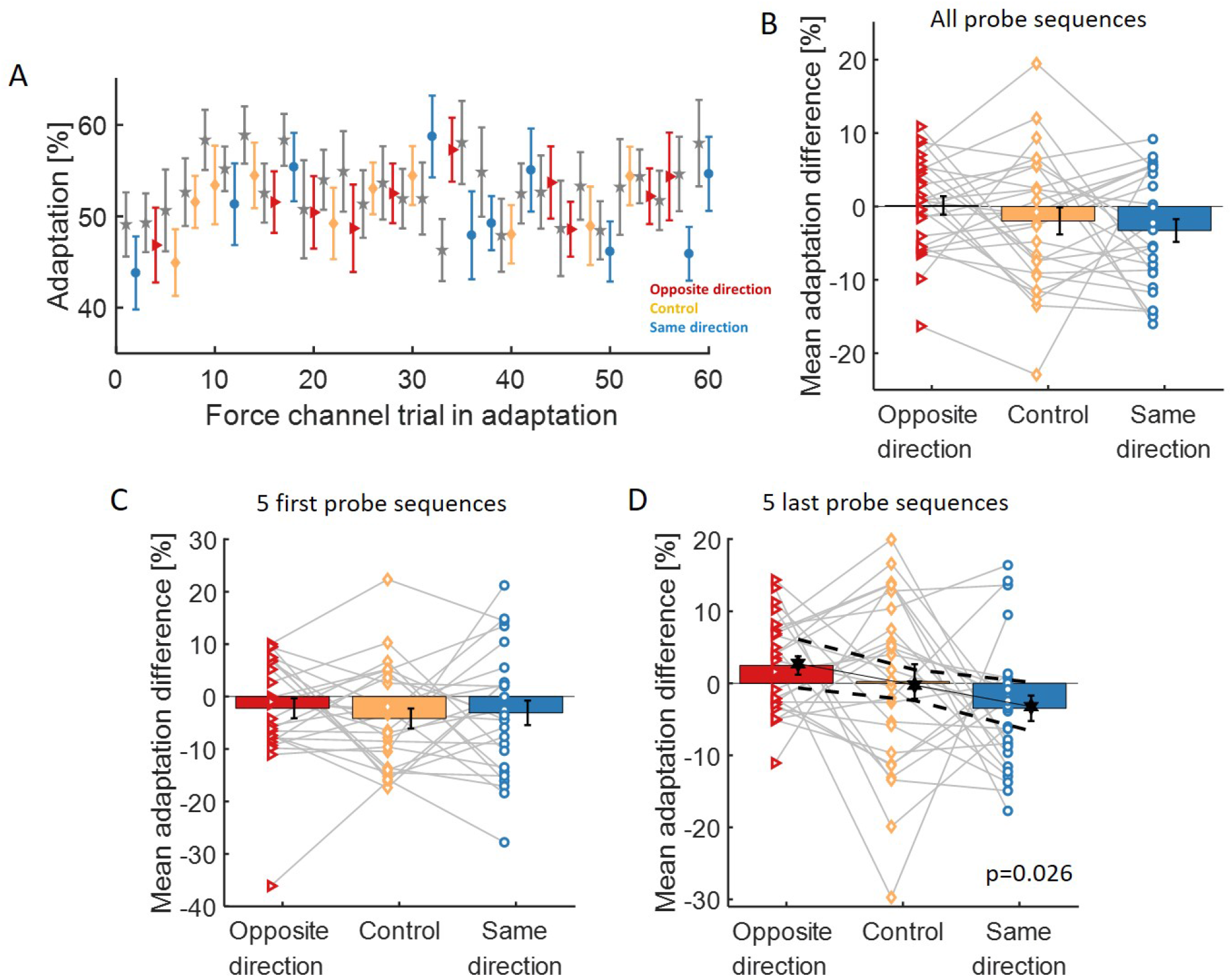
Experiment 1-adaptation coefficient. **(A)** Mean adaptation coefficient in all force channel trials. Gray and colored (red, yellow, and blue) error bars signify the first and second force channel trials within a probe sequence, respectively. **(B)** Mean difference of adaptation coefficient between the two force channel trials within all probe sequences. The markers represent the data of each participant, and the gray lines link the mean values of different gains of skin stretch for each participant. **(C)** and **(D)** are as in (B) for the five first and five last probe sequences, respectively. The solid black line represents significant regression between the three mean values, error bars are for ±SE, and the dashed black line represents ±CI.

In a post-hoc assessment, we examined whether skin stretch had a dynamic effect even after the positional error was already at a steady state. To that end, we examined whether there was a difference between the beginning and end of the exposure to skin stretch (five first, Fig. 6C, and last, Fig. 6D, probe sequences). The analysis revealed that the effect of a decreased adaptation coefficient for the same direction stimulation was consistent between the beginning and end of the exposure. However, at the beginning of adaptation, the skin stretch in the opposite direction to the force field slightly decreased the adaptation coefficient. This could be because the opposite direction of skin stretch might be confusing as it contradicts the natural stretch caused by the force field. Nevertheless, at the end of the adaptation, the opposite direction of skin stretch caused an increase in the adaptation coefficient. The ANOVA analysis did not reveal a significant difference between the groups (*F*_2,50_ = 2.46, *p* = 0.09), whereas the regression analysis yielded a significant linear regression between the mean values (*p* = 0.026). These results are in agreement with our previous study (C. Avraham et al. 2020). They demonstrate that even when participants are exposed to a relatively subtle amount of opposing tactile stimulations (compared to the consistent and continued exposure in (C. Avraham et al. 2020)), they still modify their manipulation forces when adapting to the perturbation, consistently with substantial exposure to a constant stimulation throughout the entire adaptation.

#### Grip force

To fully understand the effect of the tactile stimulation on the forces participants apply during force field adaptation, we also examined the changes in the applied grip forces during the experiment. In experiment 1, the artificial tactile information had a pronounced effect on the different components of the grip force.

##### Predictive grip force

To assess the effect of the tactile stimulation on the participants’ representation of the forces, we calculated the baseline and modulation ratio and examined the difference between the second and first force channel trials within a probe sequence (Fig. 7A-D). The baseline grip force exhibited a significant main gain effect (*F*_2,50_ = 5.21*, p* = 0.008). It increased after exposure to skin stretch in the same direction as the force field compared to no stimulation (*t*_50_ = 3.19*, p* = 0.007, Fig. 7B). A nominal increase of the baseline grip force for the opposite direction stimulation was also observed, although this effect was not statistically significant (*t*_50_ = 1.99*, p* = 0.15). These results suggest that following the same direction stimulation, participants were either insecure regarding the external environment or misinterpreted this stimulation as an increase in external force.

**Figure 7:**
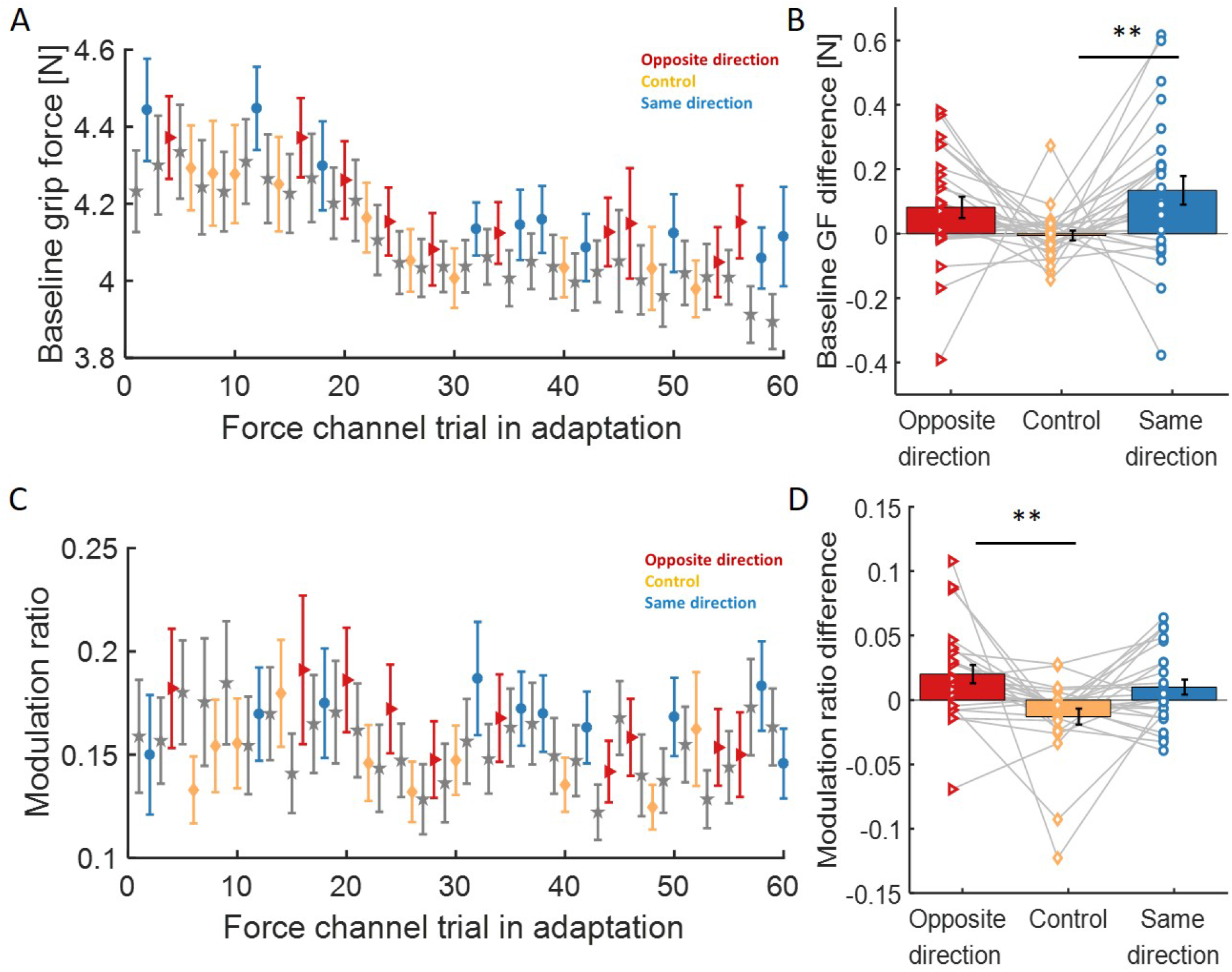
Experiment 1-Predictive grip force. **(A)** Baseline grip force in all force channel trials. Gray and colored (red, yellow, and blue) error bars signify the first and second force channel trials within a probe sequence, respectively. **(B)** Mean difference of baseline grip due to a single exposure with force field and artificial skin stretch. Markers represent each participant’s data, and the gray lines link the mean values of different gains of skin stretch for each participant. Asterisks represent significant differences: *∗ ∗ p <* 0.01. **(C)** and **(D)** are as in (A) and (B) for the modulation ratio. Error bars are for ±SE.

The modulation between the grip force and the load force was significantly affected by the skin stretch (*F*_2,50_ = 6.55*, p* = 0.003). As depicted in figure 7D, the modulation ratio increased both for the opposite direction stimulation (significant effect: *t*_50_ = 3.53*, p* = 0.003) and for the same direction stimulation (non-significant: *t*_50_ = 2.45*, p* = 0.052) compared to no stimulation. Thus, the stimulation improved the modulation between grip force and load force. Overall, the results show that the tactile stimulation affected the internal representation of the force field used for grip force control.

##### Reactive grip force

The participants’ reaction to the perturbation was examined by comparing the change in the grip force measured on a force field trial with stimulation to a prior force field trial with no stimulation (Fig. 8). This analysis revealed that for both the same and opposite direction skin stretches, the modulation increased compared to no stimulation condition (main effect: *F*_2,50_ = 21.54*, p <* 0.001, same direction: *t*_50_ = 6.53*, p <* 0.001, opposite direction: *t*_50_ = 2.68*, p* = 0.029, Fig. 8C-D). In addition, the increase of the modulation grip force was significantly higher for the same direction stimulation compared to the opposite direction stimulation (*t*_50_ = 3.85*, p* = 0.001, Fig. 8D).

**Figure 8:**
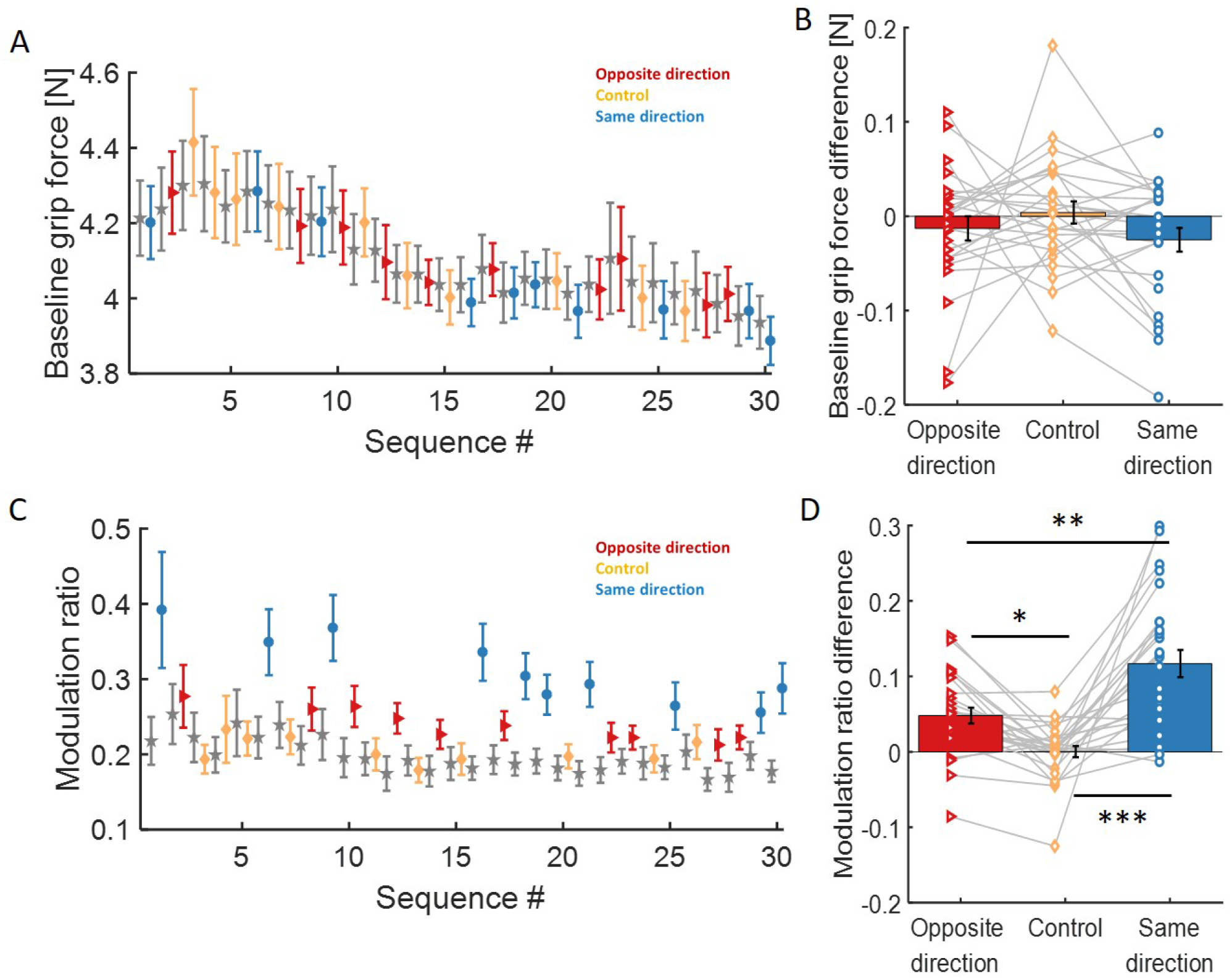
Experiment 1-Reactive grip force. **(A)** Baseline grip force. Gray and colored (red, yellow, and blue) error bars signify the force field trial proceeding and within a probe sequence, respectively. **(B)** Mean difference of baseline grip forces between the two force field trials (within the sequence minus before the sequence) over all probe sequences. Markers represent each participant’s data, and the gray lines connect the mean values of different gains of skin stretch for each participant. **(C)** and **(D)** are as in (A) and (B) for the modulation ratio. Asterisks represent significant differences: *∗p <* 0.05, *∗ ∗ p <* 0.01, *∗ ∗ ∗p <* 0.001. Error bars are for ±SE.

Overall, the grip force analysis results show an increase in both predictive and reactive components of the grip force for both same and opposite-direction stimulations compared to no skin stretch. These results are in contrast to our previous study, where we observed an increase of both the predictive and reactive grip force components only for the same direction skin stretch. This might be because here we exploited a single exposure design, which did not allow the participants to fully assess the opposite direction stimulation, prompting them to increase the applied grip force per load force amounts.

When examining these results in light of our predictions, the second hypothesis, in which we suggested that the artificial skin stretch in the opposite direction to the force field affects force field adaptation through decreased arm co-contraction, which in turn decreased the grip force, seems less likely. This is because the opposite direction stimulation increased the adaptation coefficient while the grip force also increased (instead of decreasing). An alternative explanation is that the applied stimulation increased the slippage sensation, causing the participants to increase the applied grip force to preserve a stable grip on the object.

### Augmented tactile stimulation in the same direction as a perturbing force increases force compensation

In experiment 1, we corroborated our previous finding that when the skin stretch is applied together with the force field and in the same direction, it reduces adaptation. Both models explain these findings. However, they form radically distinct predictions if the tactile stimulation is provided separately from the force field following adaptation. The haptic cuing model predicts an opposite effect between the two conditions of skin stretch with and without the force field, while the error integration model predicts a similar effect between the two conditions. Experiment 2 was designed to test the two models directly. Therefore, in experiment 2, the tactile stimulation was separated from the force field and presented during half of the force channel trials. Here, we considered three groups, each receiving a different gain of skin stretch (negative, null, or positive). To assess the effect of the skin stretch on the adaptation, we compared the kinematic and dynamic data of the three groups.

#### Directional error

We first examined the directional error during adaptation, to verify that all groups adapted to the perturbation. The results showed a similar pattern across all three groups (Fig. 9A). Namely, the directional error increased when the participants first encountered the force field, gradually decreasing throughout the adaptation. Following an abrupt removal of the force field, the participants deviated in the opposite direction, causing a sudden decrease in the directional error to negative values. We examined the difference between the different stages of the experiment (five trials from each block): the end of the Baseline block (Late Baseline), the beginning and end of the Adaptation block (Early and Late Adaptation, respectively), and the beginning of the Washout block (Early Washout). The analysis yielded a significant main effect of the stage (*F*_3,81_ = 210.13*, p <* 0.001, Fig. 9B), demonstrating that the force field did affect the trajectory during movement. When examining whether there was a difference between the groups in directional error, we found no significant group effect (*F*_2,27_ = 0.91*, p* = 0.41) and no interaction between group and stage (*F*_6,81_ = 0.87*, p* = 0.52). This indicates no difference in adaptation and aftereffects between the three groups. Additionally, the cluster permutation analysis did not reveal any clusters with a difference in directional error between the groups during the entire experiment. Thus, we conclude that the skin stretch does not affect movement trajectory during force field adaptation when applied in isolation, similarly to when the skin stretch is presented together with the force (as in (C. Avraham et al. 2020)).

**Figure 9:**
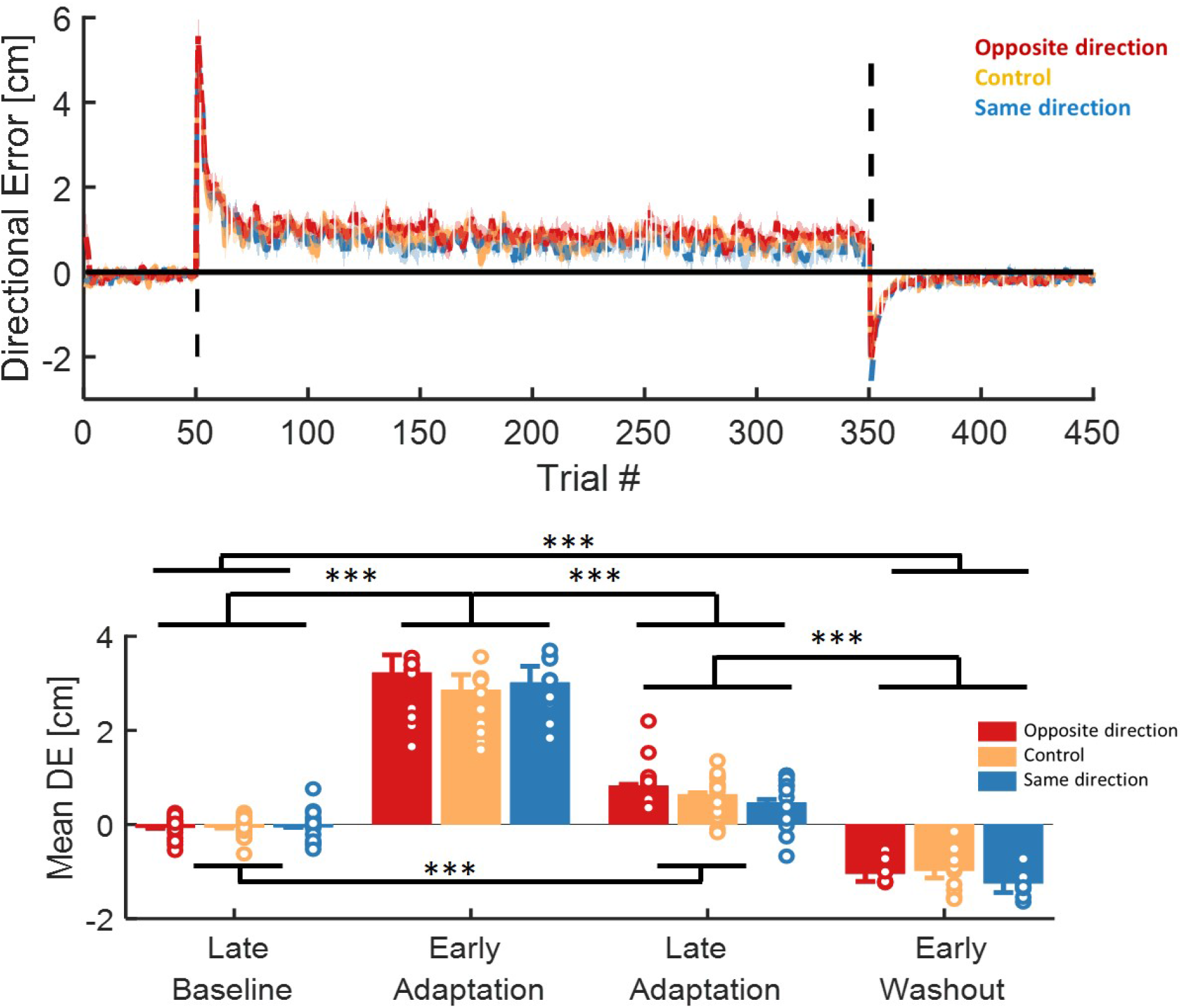
Experiment 2-Directional error: maximum deviation from a straight line. **(A)** Mean directional error across participants for each group: skin stretch in the opposite direction to the force field (red), control (no stretch, yellow), and skin stretch in the force field direction (blue). **(B)** Mean directional error of 15 trials from the different phases of adaptation: before exposure to force field (Late Baseline), the beginning of exposure (Early Adaptation), the end of exposure (Late Adaptation), and the beginning of post-exposure phase (Early Washout). Markers represent the data of individual participants. Asterisks represent significant differences: *∗∗∗p <* 0.001. Shaded regions are for ±SE.

#### Force compensation

To examine which of our models is consistent with the participants’ behavior, we analyzed the force compensation during force channel trials and compared the adaptation coefficient between the three groups.

Strikingly, we found an opposite pattern to the one found in Experiment 1. I.e., the adaptation coefficient was higher for the same direction groups than for the opposite direction group. This pattern was evident both in force channel trials with (Fig. 10B) and without (Fig. 10A) stimulation. When looking for clusters of interest throughout the entire adaptation period, the analysis did not reveal any clusters of interest. However, when we examined the difference between the beginning and end of the adaptation, the ANOVA analysis yielded a significant main effect of the stage (without stimulation: *F*_1,42_ = 92.589*, p <* 0.001, Fig. 10C, with stimulation: *F*_1,42_ = 171.48*, p <* 0.001, Fig. 10D), confirming that the participants were able the adapt to the force field in all groups. Critically, the statistical analysis also yielded a significant group X stage interaction effect only in force channels with stimulation (without stimulation: *f*_2,42_ = 1.89*, p* = 0.168, with stimulation: *f*_2,42_ = 3.25*, p* = 0.048, Fig. 10D). A post hoc t-test revealed a significant difference between the two groups of the same and opposite direction skin stretch (*t*_42_ = 2.1*, p* = 0.04). Accordingly, we conclude that when the artificial information is congruent with the direction of the external force field and applied in force channel trials, the artificial information improves the force field learning and vice versa.

**Figure 10:**
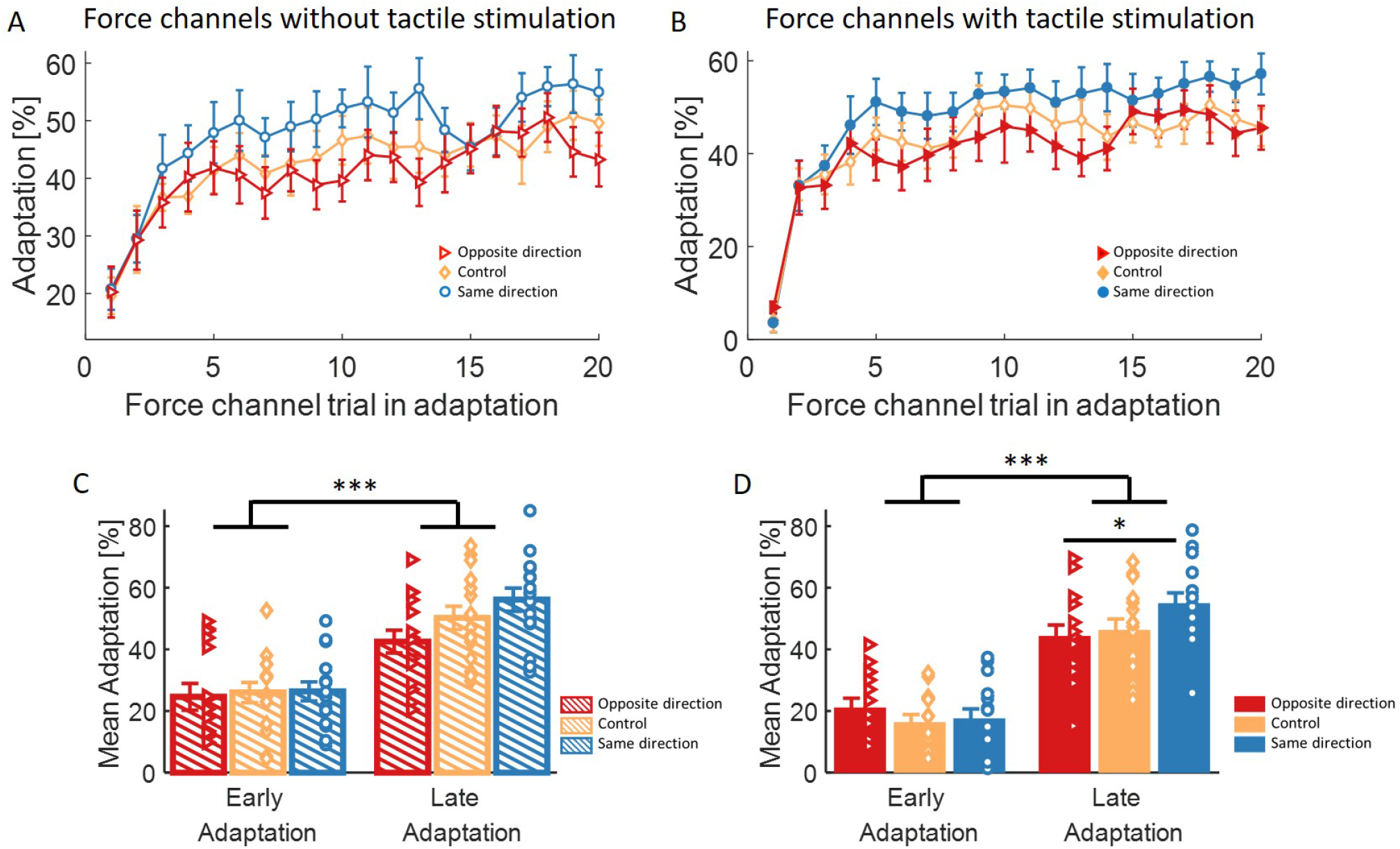
Experiment 2-Adaptation coefficient. **(A)** Mean adaptation coefficient measured in force channel trials without the stimulation for the group with skin stretch in the opposite direction to the force field (red), control (no stretch, yellow), and force field direction (blue). **(B)** Same as (A) for force channel trials with the skin stretch in the same or opposite direction to the force field (blue and red filled markers, respectively), and force channel trials without stimulation for the control group (empty yellow markers). **(C)** Mean adaptation over two first force channel trials (early) and two last force channel trials (late) without tactile augmentation. Markers represent the data of each individual subject. **(D)** Same as (C) for force channel trials with tactile augmentation. Asterisks represent significant differences: *∗p <* 0.05, *∗ ∗ ∗p <* 0.001. Error bars are for ±SE.

The results of experiment 2 are consistent with the haptic cuing model (Fig. 4 left panel) and stand in stark contrast to the prediction of the error integration model. This suggests that tactile stimulation is used by the sensorimotor system as a guidance cue, directly affecting the motor output during the learning process.

#### Grip force

In experiment 1, we found that when the tactile stimulation is applied together with the force field, there is an effect on the applied compensation forces as well as on the grip forces applied to stabilize the device during the movement. To examine the effect of the skin stretch that is separated from the force field on the applied grip forces, we calculated the baseline grip force and modulation ratio and compared the results between the three groups. We found an increase in the modulation grip force at the beginning of adaptation for the group with skin stretching in the same direction as the force field (Fig. 11A-C). This increase was observed mainly in force channel trials with the stimulation (Fig. 11B). The ANOVA analysis did not show a difference between the beginning and end of adaptation. However, when examining the entire adaptation period, we saw clusters with a significant difference between the same direction group and both opposite and control groups. Moreover, this increase was also observed in force channel trials without stimulation (Fig. 11C) and force field trials (Fig. 11A, although insignificant). To examine whether this increase during the beginning of adaptation was due to a predictive or reactive effect, we examined whether there was a difference between the modulation ratio in force channel trials with the stimulation (representing both predictive and reactive components) compared to force channel trials without the stimulus (representing only the predictive component, Fig. 11D). We found no stages and clusters with a significant difference between the two types of trials. Therefore, we claim that the increase in the grip force is mainly due to the predictive modulation component of the grip force. In the case of the baseline grip force, there was also an increase for the group with skin stretch in the force field direction (Fig. 12A-C). However, both ANOVA and cluster analysis showed no significant differences between the groups. Nevertheless, the observed increase in both modulation and baseline grip forces might be due to a slippage sensation, which causes an increase in the applied grip force to prevent the object from slipping.

**Figure 11:**
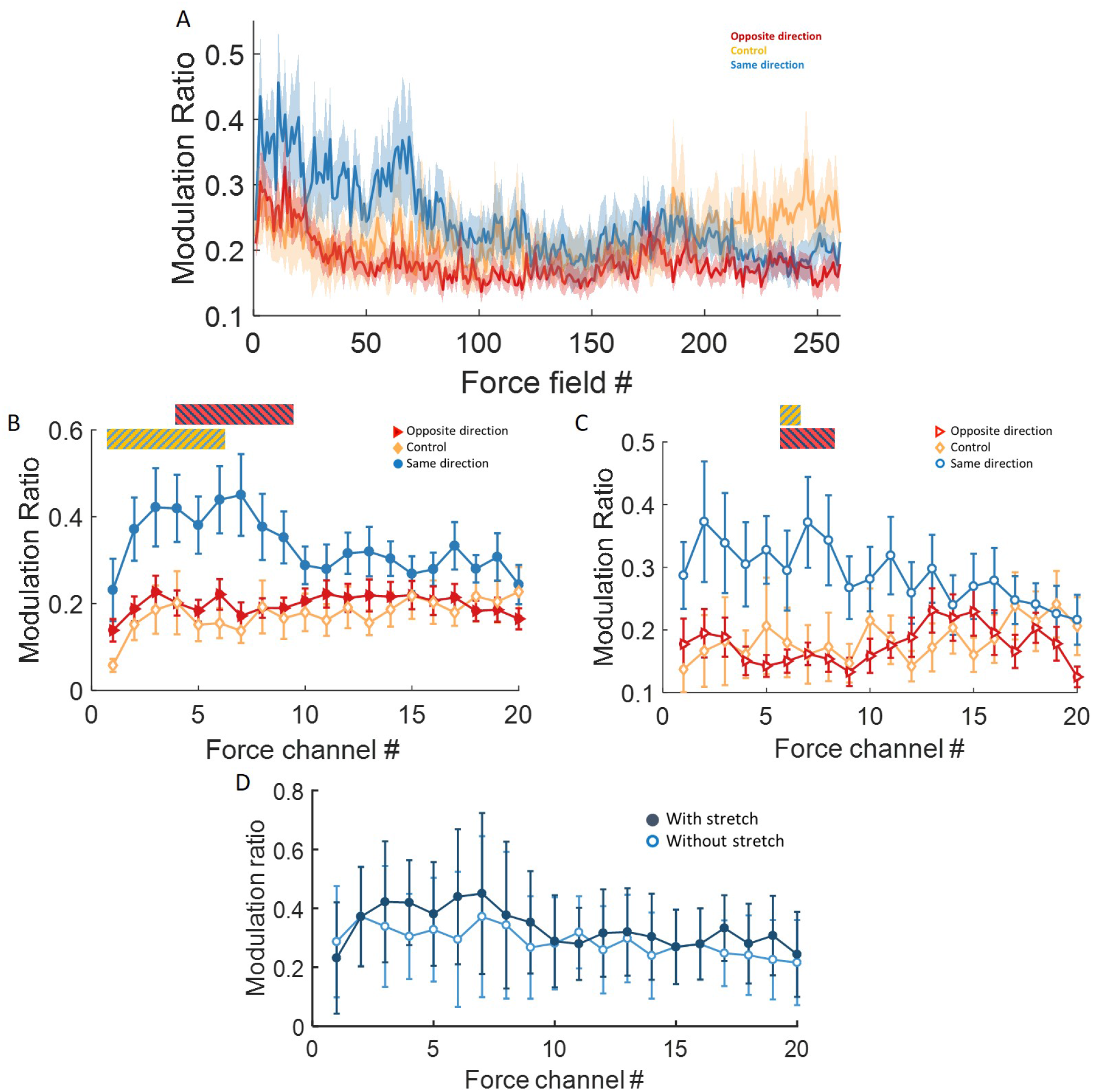
Experiment 2-Modulation ratio: the modulation of the grip force according to the external load force. **(A)** Modulation ratio in all force field trials for the three groups of skin stretch in the opposite direction to the force field (red), control (no stretch, yellow), and force field direction (blue). The solid line represents the mean value. **(B)** Modulation ratio in all force channel trials accompanied by augmented tactile stimulation. Filled lines with stripes represent clusters that are statistically different between the two groups. Blue and red stripes indicate significant differences between the same and opposite direction groups, respectively, while yellow and blue stripes indicate significant differences between the same direction group and the control group, respectively. **(C)** Modulation ratio in all force channel trials without tactile stimulation. Colors, lines, and markers are as in (A) and (B). **(D)**Modulation ratio in force channel trials with (dark blue) and without (light blue) skin stretch for the group with skin stretch in the same direction of the force field. Shaded region and error bars are for ±SE.

**Figure 12:**
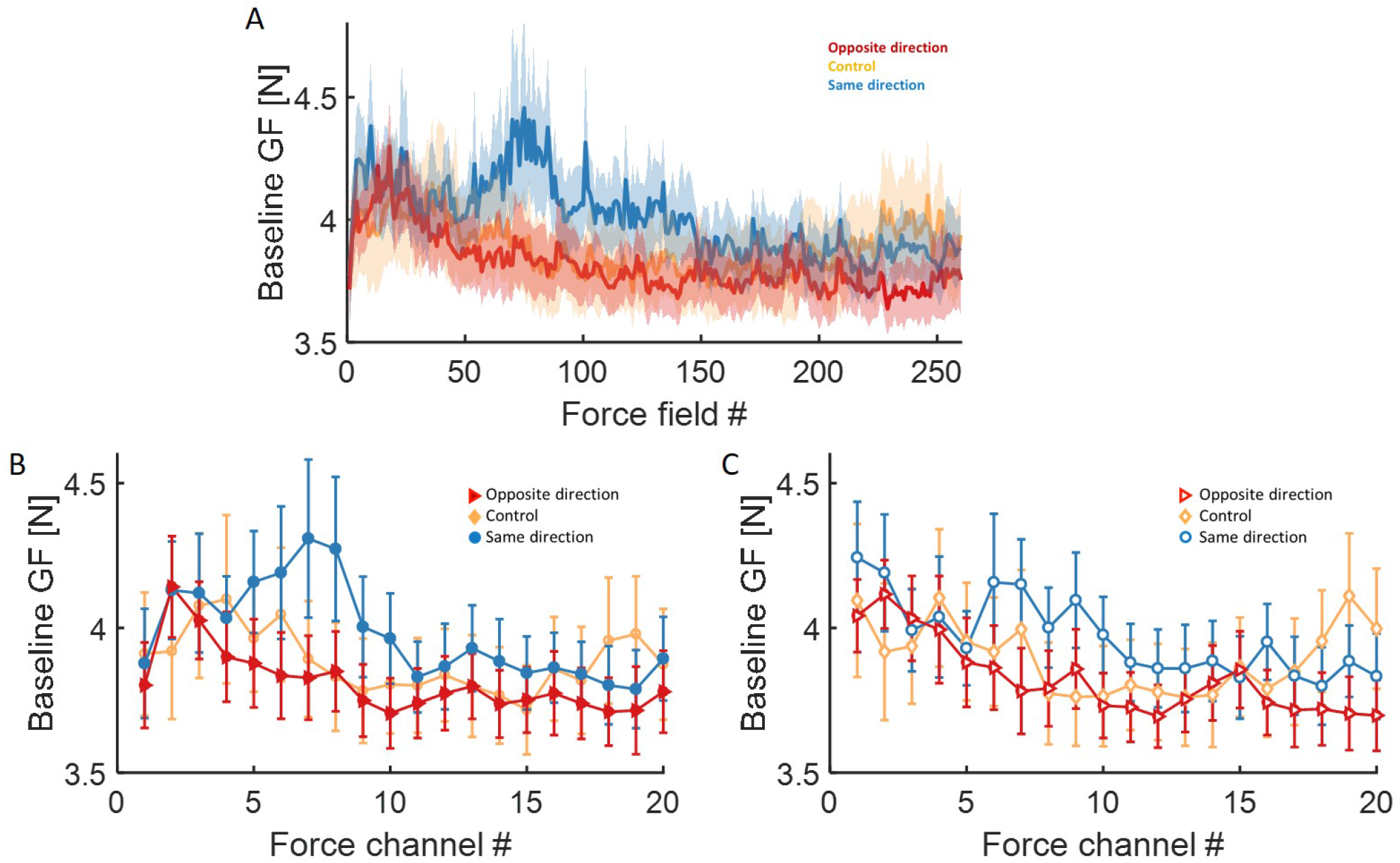
Experiment 2-Baseline grip force analysis. **(A)** Baseline grip force in all force field trials. We present the grip force at the beginning of the movement (t=0) for the three groups of skin stretch in the opposite direction to the force field (red), control (no stretch, yellow), and the force field direction (blue). The solid line represents the mean value. **(B)** Baseline grip force in all force channel trials accompanied by augmented tactile stimulation. Colors are as in (A). **(C)**Baseline grip force in all force channel trials without tactile stimulation. Shaded region and error bars are for ±SE.

Overall, the grip force analysis in experiment 2 showed that the skin stretch in the force field direction modified the internal representation used for grip force control, increasing the predictive component of the grip force. This further supports the notion that our second hypothesis, suggesting the tactile stimulation affected the force field adaptation through arm co-contraction is less likely. This is because, according to this hypothesis, the compensation and grip forces are updated via a similar mechanism. In contrast, when comparing the two experiments, we can see the opposite effect on the compensation forces, while a similar effect is observed on the grip force.

Evidence from our modeling work and the results from both experiments strongly support the first hypothesis, according to which tactile stimulation affects the compensation and grip forces differently. The compensation forces are updated through guidance cuing, while the grip forces are affected by a separate mechanism, possibly via a slippage sensation. Additional support for this claim is provided by our EMG analysis, which shows a similar effect on arm co-contraction between the two experiments, suggesting that it was not the mechanism through which tactile stimulation affected the adaptation to the force field (see supplementary material).

## Discussion

In this study, we show that artificial tactile stimulation affects adaptation to novel dynamics, both when the stimulus is congruent or incongruent with the direction of the force field and when the stimulus is added during force field or force channel trials. Consistent with our previous results (C. Avraham et al. 2020), skin stretch stimulation applied simultaneously with the force field reduced compensatory forces when occurring in the same direction of the force, while the adaptive response increased when it was in the opposite direction. Strikingly, when the skin stretch was provided during force channel trials and separately from the force field that drove adaptation, it led to an opposite pattern of effect: the skin stretch in the same direction increased compensatory forces, whereas skin stretch in the opposite direction attenuated them. Our modeling work suggests these results are consistent with the idea that artificial skin stretch is used as a guidance cue for the “appropriate” motor command rather than interpreted as an error signal integrated into the overall force prediction error.

Force field adaptation has been investigated extensively over the years (Conditt et al. 1997; R. Shadmehr et al. 1994; Scheidt et al. 2000; Yousif et al. 2012; Gonzalez Castro et al. 2014; Schween et al. 2020). It has been recently suggested that such an adaptation process is driven by both implicit (unconscious) and explicit (deliberate) learning systems (Schween et al. 2020). Key distinctions between the two systems manifest in the rate at which they develop; implicit learning develops slowly, while explicit learning is faster (Krakauer et al. 2019; Kim et al. 2021). In experiment 2, we show an effect of the skin stretch on adaptation but no difference in the aftereffects between the three groups, which are a measure of implicit learning (Taylor et al. 2014). In addition, modeling the contribution of the skin stretch to adaptation revealed that the model best fitting our results was the one where the skin stretch directly affected the motor output with no changes to the representation of the force error. This effect on the motor output can be viewed as a guidance cue affecting explicit strategies without altering the implicit representation of the external forces. Consistent with this view, the result of experiment 1 suggests that although the participants were exposed to two opposite tactile stimuli, there was still an effect of adaptation rather than interference. This suggests that the effect of the skin stretch on adaptation can be switched off relatively easily – a property that characterizes explicit learning (Morehead et al. 2015; Vandevoorde et al. 2019; G. Avraham et al. 2021). Therefore, we speculate that the skin stretch explicitly affects the adaptation by modifying the motor output in each trial. It will be interesting to further examine this assumption in future studies by developing an experimental protocol to directly assess the effect of tactile stimulation on implicit and explicit learning.

In this study, we show evidence for different mechanisms that update the manipulation and grip forces during adaptation. This is demonstrated by an opposite effect of the tactile stimulation on the manipulation forces between the two experiments, yet a consistent effect on the grip force. Previous motor learning studies showed a dissociation between manipulation and trajectory control to grip force control. Namely, it has been demonstrated that grip force is modulated in the early stages of adaptation solely by the experienced kinetic errors (J. Randall Flanagan et al. 2003; Danion et al. 2013). In addition, to prevent slippage, a safety margin is added to the grip force, proportional to the variability of the load force (Hadjiosif et al. 2015). In contrast, the adaptation of manipulation forces and trajectory control is developed more slowly and driven primarily by kinematic errors (J. Randall Flanagan et al. 2003; Danion et al. 2013), being sensitive to the average value of the load force (Scheidt et al. 2001; Mawase et al. 2012; Hadjiosif et al. 2015). Moreover, in the context of tactile stimulation, studies showed that artificial tactile stimulation had no effect on the performed trajectory during adaptation (Rosati et al. 2014; C. Avraham et al. 2020), but a rather pronounced effect was observed on the grip force control (C. Avraham et al. 2020; Farajian et al. 2020). Therefore, our results support the idea that manipulation and trajectory control are updated through different mechanisms than the ones used for grip force control.

In motor learning, multiple input signals can be used as cues that improve learning. Namely, studies have examined visual cues (Addou et al. 2011), auditory cues (Schaefer 2014), and haptic cues (Crespo et al. 2008; Grindlay 2008). In the case of providing a directional cue during force field adaptation, it has been shown that upon receiving a visual cue that is congruent with the direction of the force field, participants exhibit a higher adaptation level, compared to participants that were provided with incongruent or constant visual cues (Franklin et al. 2023). Compared to visual cues, haptic cues are potentially more salient in improving motor learning, especially in adaptation to the force field, where the task itself is haptic in nature. However, studies have shown contradicting results regarding its benefit in promoting motor learning in terms of retaining motor skills after learning (Lee et al. 2010). In the same context, it was also shown that recalling explicit strategies enhanced savings of previously learned motor processes (Morehead et al. 2015; Haith et al. 2015; G. Avraham et al. 2021). Here, we show that tactile haptic feedback might be interpreted differently, as a guidance cue, depending on whether it is provided with or without the perturbing force, affecting the extent of adaptation. Nevertheless, an interesting question that our study raises is whether this guidance effect, which can be considered explicit, is retained upon re-exposure to the same force field and whether it would promote the recall of strategies that have been previously proven useful.

An important aspect that might contribute to the results of our first experiment is the Simon-left effect, according to which the spatial location of a stimulus will affect one’s response accuracy, such that when the stimulus and the response are on the same side – the response will mostly be faster and more accurate (Hommel 1993). In our case, when the stimulus was in the same direction as the participants’ manipulation forces, adaptation was improved. However, this effect can not explain the results of our second experiment, in which adaptation was improved when the skin stretch was in the opposite direction to the required manipulation forces. Nevertheless, in this experiment, the stimulus and response were presented in separate trials. To assess whether part of the results observed in the first experiment could be attributed to the Simon-left effect, it will be interesting to examine force field adaptation wherein the force field and tactile stimulation are provided in forward and backward directions, while the movements are lateral. In this paradigm, we will eliminate the side effects while preserving the relation between stimulus and force direction.

Our results point to a large inter-participant variability in the effect of skin stretch on force field adaptation. This is consistent with numerous skin stretch stimulation studies in which participants exhibited large variability in perceptual effects (Farajian et al. 2020; Z. F. Quek et al. 2013; C. Avraham et al. 2020; Farajian et al. 2021; Kossowsky et al. 2022). This variability might stem from the amount of skin stretch being dependent on various properties of the skin, which is known to change dramatically with age and environmental conditions (Yang et al. 2018; Langton et al. 2017; Deflorio et al. 2022). In addition, the sensitivity to tactile feedback can vary with device and feedback characteristics (Valle et al. 2018; Jing et al. 2021). In addition, in this study, the sensitivity might have changed during the construction of the internal representation when adapting to the new dynamics. To address several of these issues, it will be interesting to measure the amount of skin deformation during the adaptation process (Delhaye et al. 2021; Willemet et al. 2021), and thereby assess the causal relationship between the amount of stretch and manipulation force control extent.

Tactile stimulation gained considerable interest in the past few years. This stimulation can be delivered to the skin using vibration (Yatani et al. 2009), pressure (Aggravi et al. 2018), or tangential deformation (Gleeson et al. 2010); it can thereby convey multiple types of information, such as position (Bark et al. 2008), force augmentation (Porquis et al. 2011), or friction (Provancher et al. 2009). In our experiment, a clear influence of the stimulation’s direction is observed. Therefore, it is safe to assume that this stimulation was not perceived as noise that can be ignored but rather as an information signal that should be considered during adaptation. Moreover, we rule out the possibility that the effect of skin stretch is attributed to misinterpretation of the external forces’ magnitude (using our model) or to muscle co-contraction. That being said, our results suggest that when the tactile stimulation is applied in a direction that can be interpreted as an assistive cue (such as guidance or force cue), it will improve the adaptation. Otherwise, the stimulation will cause confusion or an increase in the cognitive load that leads to impaired adaptation. These results strongly support that kinesthetic and tactile information are integrated during sensorimotor learning in a way that appears to be complicated and task-related.

From the extensive research on the effect of artificial tactile information, it is clear that this stimulation can be exploited to elicit many types of sensations and, thus, can be used for various applications. If the additional stimulation is presented together with the dynamics, skin stretch in the opposite direction from the external dynamics will improve adaptation. If a certain scenario requires performing movements without force information about the external dynamics, stretch with the same properties as the external dynamics can improve adaptation. Using these results for future technologies, tactile stimulation can enhance motor learning and improve the efficacy of assistive and rehabilitation haptic devices.

## Supplementary material

To better understand the effect of artificial tactile information on force field adaptation, we examined whether differences in muscle co-contraction can explain the observed effect on the adaptation coefficient. This is to verify that our second hypothesis, suggesting an effect of the tactile stimulation on arm co-contraction, is not valid. Therefore, we examined the integrated EMG (iEMG) as a measure of increased muscle activity. We separated between agonist and antagonist pairs of muscles (flexors: Deltoid Lateral, Biceps Short, Biceps Long, Flexor Carpi Radialis, and Flexor Carpi Ulnaris. Extensors: Triceps Long, Triceps Lateral, and Extensor Carpi Ulnaris), and examined the average effect over each group.

For the flexor muscles group, we saw a non-significant trend in the effect on muscle activity. In experiment 1, we found no difference between the iEMG in the two force channel trials within a probe sequence (Fig. S1A). However, when we compared the two force field trials with and without the skin stretch (i.e., the force field trial within and before the probe sequence, Fig. S1B), we found a trend for an increase in the iEMG value after receiving the tactile stimulation in the force field direction. Moreover, a slight decrease was observed for the opposite-direction stimulation. Although these results were not statistically significant (ANOVA: *F*_2,50_ = 1.62, *p* = 0.21, regression analysis: *p* = 0.163), they show a trend in muscle activity during the experiment. In experiment 2, there was also an increase of the iEMG for the group with skin stretch in the force field direction compared to the opposite direction group. This effect was also not significant due to the large variability (ANOVA: *F*_2,42_ = 1.3, *p* = 0.28, Fig. S1C). Accordingly, in both experiments, the trend in muscle co-contraction is similar, even though an opposite effect was observed in the manipulation forces results. Hence, we claim that the skin stretch effect on adaptation was not caused by muscle co-contraction.

**Figure S1:**
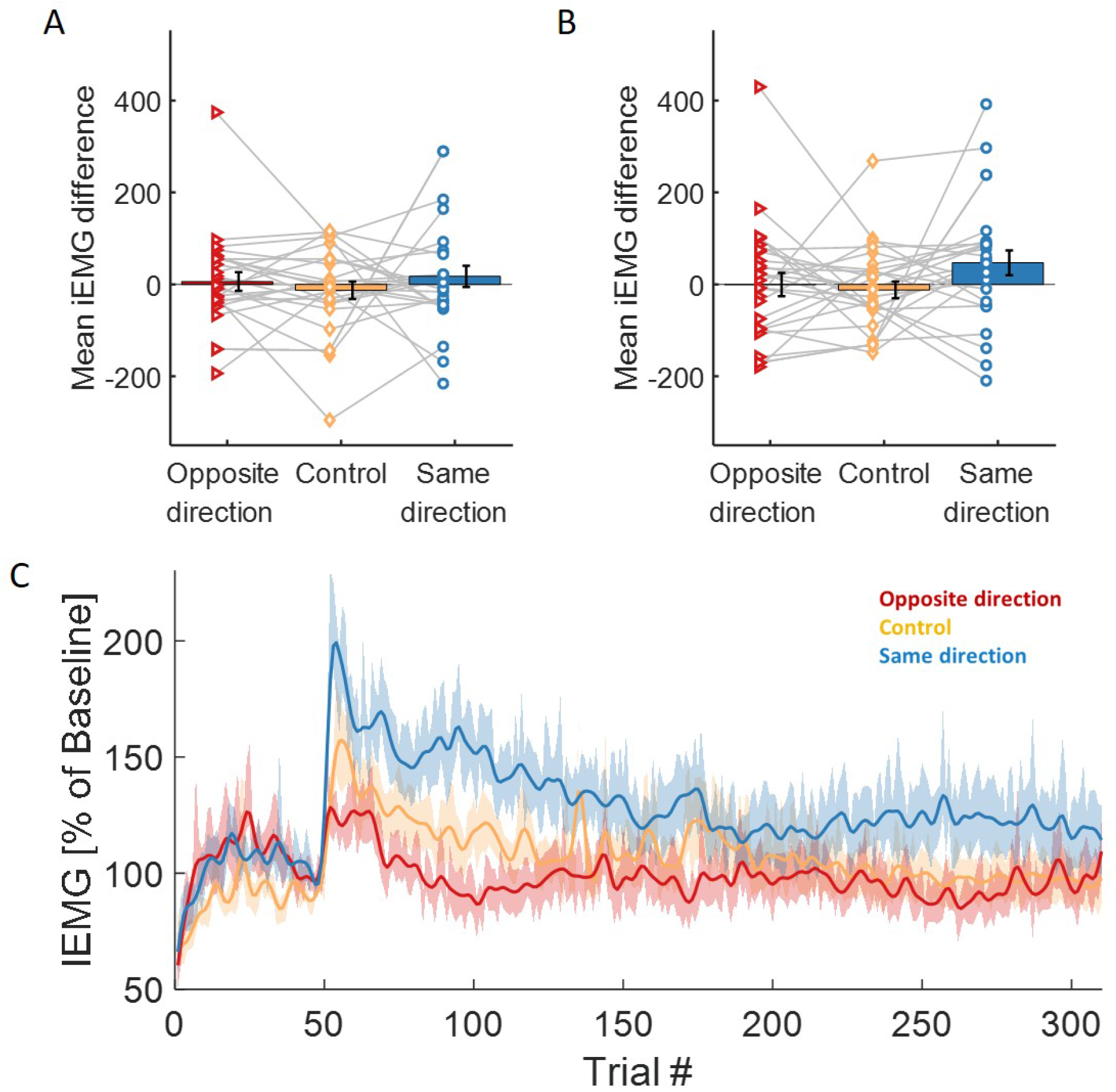
EMG analysis. **(A)** Mean difference of integrated EMG measured between the two force channel trials within a probe sequence (second minus first) in experiment 1 for the skin stretch gains (red, yellow, and blue for g=-100,0,100, respectively). Markers represent each participant’s data, and the gray lines connect the mean values of different gains of skin stretch for each participant. **(B)** Mean difference of integrated EMG between the force field trial within the probe sequence to the force field trial that proceeds the probe sequence. Colors, markers, lines, and error bars are as in (A). **(C)**Integrated EMG in the Baseline and Adaptation blocks (excluding force channel trials) obtained in experiment 2. For each participant, we normalized the iEMG values at the Baseline and Adaptation blocks by the iEMG in the Baseline block (5 last trials). The solid line represents the mean value. Here, the colors mark the three groups in the experiment of skin stretch in the opposite direction to the force field (red), control (no stretch, yellow), and the force field direction (blue). Shaded region and error bars are for ±SE.

## Acknowledgments

The authors wish to thank Dr. Raz Leib for his assistance with analyzing the EMG signals. The study is supported by the Binational United-States Israel Science Foundation (grant no. 2016850), the National Science Foundation (grant no. 1632259), the Israeli Science Foundation (grant 823/15), and the Helmsley Charitable Trust through the Agricultural, Biological, and Cognitive Robotics Initiative of Ben-Gurion University of Negev, Israel. CA was supported by the Besor Fellowship and the PBC scholarship.

## Contribution

CA, GA, and IN designed the experimental protocol and hypotheses. CA performed the experiments. CA analyzed the data. CA, GA, and IN interpreted the results, and CA wrote the first draft of the paper. CA, GA, and IN edited the paper and approved the final version.

## Data availability

The Matlab code for the state-based models and the SolidWorks parts of the skin-stretch device will be provided upon request from the corresponding author.

